# Alpha-1 Antitrypsin Overexpressing Mesenchymal Stem/Stromal Cells Reverses Type 1 Diabetes via Promoting Treg Function and CD8^+^ T cell exhaustion

**DOI:** 10.1101/2025.04.19.649667

**Authors:** Hua Wei, Wenyu Gou, Judong Kim, Suganya Subramanian, Tiffany Yeung, Paramita Chakraborty, Shikhar Mehrotra, Stefano Berto, Charlie Strange, Hongjun Wang

## Abstract

Mesenchymal stem/stromal cell (MSC) therapy holds great promise as both a therapeutic option and as a biofactory, as cells produce therapeutic proteins to augment their efficacy in disease treatment. This study investigates the therapeutic effects and the mechanistic insights of alpha-1 antitrypsin overexpressing MSCs (AAT-MSCs) in diabetes prevention and treatment. A single infusion of AAT-MSCs not only delayed diabetes onset but reversed new-onset type 1 diabetes (T1D) in the nonobese diabetic (NOD) mice. Using single-cell RNA sequencing, flow cytometry, and functional analyses, we characterized the impact of AAT-MSCs on immune cells, particularly CD4^+^ and CD8^+^ T cells, in pancreatic lymph nodes (PLNs) and islets of NOD mice. AAT-MSCs enhanced the immunosuppressive function and the communication of regulatory T cells (Tregs) with other immune cells while reducing the numbers of T helper 1 (Th1) cells and CD8^+^ cytotoxic T cells. *In vitro* experiments further confirmed the capacity of AAT-MSCs to promote the proliferation of Tregs, which consequently fostered an exhausted phenotype in CD8^+^ T cells, thereby facilitating β cell survival and potentially aiding in diabetes remission. Thus, our findings underscore the significant protective effects of AAT-MSCs, delineate their novel mechanistic insight on recipient immune cells, and provide evidence for the clinical application of AAT-MSCs in treating T1D.

## INTRODUCTION

Type 1 diabetes (T1D) is characterized by autoimmune-mediated destruction of insulin-secreting pancreatic β cells that leads to hyperglycemia. Patients need life-long insulin therapy to maintain normoglycemia. Currently, limited approaches are available to delay the onset or achieve a cure for T1D ^1^. While insulin replacement therapy can effectively control glucose levels, it cannot restore pancreas function and is associated with risks such as hypoglycemia and infection ^2^. The ideal therapeutic approach for T1D would effectively restore immune system homeostasis and protect pancreatic β cells from further destruction. Currently, no single intervention can provide both of these benefits. Thus, there is a critical need to develop more effective strategies or technologies for delaying and treating T1D.

Mesenchymal stem/stromal cells (MSCs) are adult stem cells that can be isolated from various body tissues and are increasingly recognized as a promising source for cell therapy due to their immunomodulatory, tissue repair, and regenerative functions ^3^. MSCs exert protective effects by secreting pro-mitotic, anti-apoptotic, anti-inflammatory, and immunomodulatory factors while mitigating metabolomic and oxidative stress imbalances and restoring homeostasis. The immunomodulation function of MSCs has been observed in many disease models, including encephalomyelitis ^4^, T1D ^5^, multiple sclerosis ^6,7^, graft-versus-host disease (GvHD) ^8^, arthritis ^9^, and others. The immunosuppressive properties are partly based on their nitric oxide production, indoleamine 2,3 dioxygenase (IDO), and transforming growth factor β (TGF-β) with complex interactions in other immune pathways ^10–14^. Data from various studies, including ours, show that systemic infusion of MSCs in murine models of diabetes improved glycemic control, reduced pancreatic insulitis, prevented autoimmune destruction, and promoted repair of pancreatic tissue ^15–18^. Importantly, intravenously infused MSCs migrate to the injured pancreas or pancreatic islets in the T1D mouse models ^19,20^. Two recent clinical trials demonstrated the preliminary safety and efficacy of MSCs in treating new-onset T1D in patients ^21,22^. The U.S. Food and Drug Administration (FDA) has recently approved Remestemcel-L (Ryoncil, Mesoblast, Inc.), an allogeneic bone marrow-derived MSC therapy, for treating steroid-refractory GvHD in pediatric patients.

In addition to their therapeutic effects, MSCs are also one of the most promising stem cell populations for use in gene therapy studies and trials, as they can be modified with a wide range of both viral and non-viral vector systems to produce therapeutic proteins, which further enhance their natural abilities to mediate repair within various tissues ^23^. Alpha-1 antitrypsin (AAT) is an acute phase reactant and serine protease inhibitor that suppresses various enzymes, including neutrophil elastase, cathepsin G, and others ^24^. It also exerts anti-inflammatory and anti-apoptotic effects via suppressing cytokine production, complement activation, and immune cell infiltration ^25^ while protecting pancreatic β cells from apoptosis ^26–29^. In non-obese diabetic (NOD) mice, a single injection of AAT reduced the intensity of insulitis, increased β cell mass, promoted β cell regeneration, and prevented the onset of diabetes via modulating Tregs ^28,30^. AAT has been shown to protect mouse and non-human primate islet grafts from failure by reducing islet cell death and promoting graft revascularization ^29,31,32,24,33^. In our previous studies, human AAT-overexpressing MSCs (AAT-MSCs) demonstrated enhanced innate properties, including increased proliferative capacity and accelerated migration ^34^. Infusion of AAT-MSCs showed better efficacy in preventing the onset of T1D in the NOD mice than control MSCs ^34,35^ and significantly better protection in GvHD^36^. This study aims to decipher the mechanistic basis of AAT-MSC action by examining their impact on immune cells, particularly Tregs and exhausted CD8^+^ T cells, at single-cell resolution in both *in vitro* and *in vivo* models, providing mechanistic insights for potential clinical applications.

## RESULTS

### 1. Treatment with AAT-MSCs delays the onset of diabetes in female NOD mice

We previously demonstrated that infusion of human bone marrow-derived AAT-MSCs delays the onset of T1D ^34^. This data was further validated by a single dose infusion of human bone marrow-derived AAT-MSCs to female NOD mice at eight weeks of age, which resulted in a substantial delay in diabetes onset compared to non-treated control mice followed up to 25 weeks of age (n=34 in non-treated control (CTR, n=32 in AAT-MSC group, **Fig. 1a-c** and ^34^). At three weeks post-infusion (11 weeks of age), a time estimated to be sufficient for MSCs to achieve protection, we measured islet function in mice using the intravenous glucose tolerance test (IVGTT). AAT-MSC-treated NOD mice exhibited consistently lower average blood glucose levels at multiple time points during the IVGTT, as well as the reduced area under the curve (AUC), compared to the control group (n=12 in AAT-MSC, n=8 in CTR, **Fig. 1d &e**). Additionally, these mice showed significantly higher basal C-peptide levels during the IVGTT, a trend of higher C-peptide AUC (**Fig. 1f &g**), and a significantly higher mixed meal stimulation index (MMSI, calculated as the C-peptide AUC, divided by the glucose AUC, **Fig 1e**), indicating improved islet function compared to controls. Insulitis, characterized by immune cell infiltration into pancreatic islets, is a hallmark of T1D. Scoring of islets based on the extent of insulitis showed that AAT-MSC-treated mice had more intact (islets with < 5% immune cell infiltration) and fewer severely infiltrated islets (islets with >75% immune cell infiltration) compared to controls (n=6 per group, (**Supplemental Fig. S1a** and **Fig. 1i&j**). Furthermore, islets isolated from AAT-MSC-treated NOD mice demonstrated a trend toward increased mRNA expressions of mature β cell markers, including *Pdx1*, *Ins1*, *Mafa*, and *NKx6.1*, compared to islets from control mice (**Supplemental Fig. S1, b-e).** These results suggest that administering AAT-MSCs at week 8 effectively delayed the onset of T1D by mitigating insulitis and preserving islet survival and/or function.

**Fig. 1.**
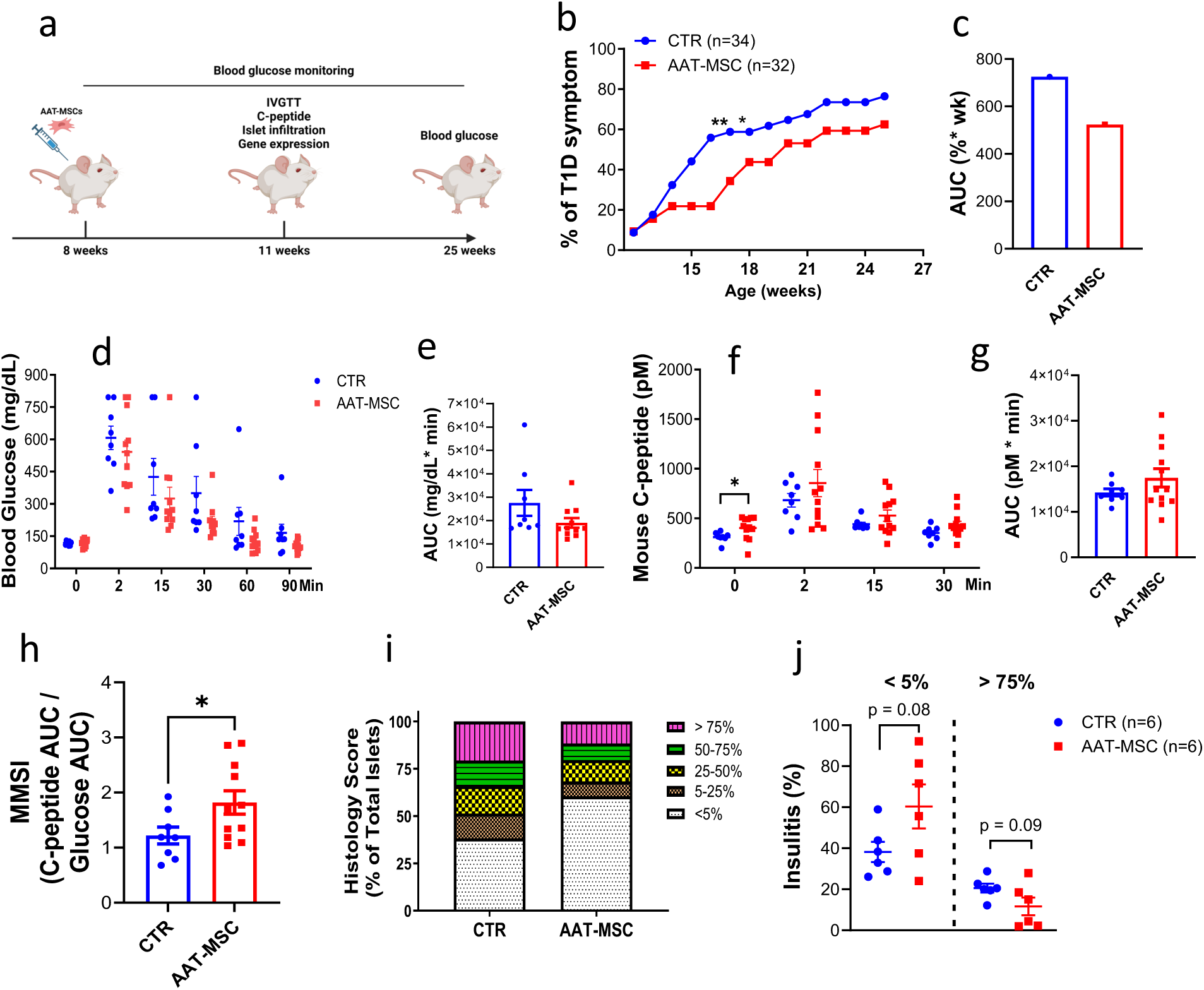
AAT-MSC infusion delays the onset of T1D in female NOD mice. **a**. Schematic diagram of the treatment protocol, sample collection, and analysis for control and AAT-MSC-treated mice. **b**. Percentage of mice with T1D onset and **c**. AUC for mice receiving AAT-MSCs (AAT-MSC, n=32) or controls (CTR, n=34). ** p < 0.01, * p < 0.05, AAT-MSC vs. CTR. P = 0.122, by log-rank test. The Z-test evaluated the difference of % mice escaping normoglycemia each week. Z value was 2.82, revealing p < 0.01 at week 16; also, Z=1.99 and p < 0.05 at week 17. AUC: area under the curve. **d.** Blood glucose levels and AUC during the IVGTT in CTR (n=8) and AAT-MSC (n=12) mice at age 11 (3 weeks post-treatment). C-peptide level **(f),** AUC of C-peptide levels **(g)** and MMSI (**h**) during IVGTT in CTR and AAT-MSC-treated mice. Each dot represents an individual mouse. *p < 0.05, Student’s t-test. **i:** Insulitis scores in CTR and AAT-MSC treated pancreatic islets. **J**. Percentage of islets with <5% or >75% of immune cell infiltration in CTR and AAT-MSC-treated islets. Each group included six mice, and individual histology scores were determined from 385 islets in the control group and 386 islets in the AAT-MSC-treated group.

### 2. Single-cell RNA sequencing (ScRNAseq) analysis reveals immune cell subtypes impacted by AAT-MSC therapy in both PLNs and islets of treated mice

The activation of immune cells in T1D begins within the islets and is amplified in lymphoid tissues, particularly in the pancreatic draining lymph nodes, which serve as a hub of aberrant immune response ^37^. To assess the effects of AAT-MSCs on the immune cell phenotypes and gene expression at single cell level, we collected PLNs or islets from AAT-MSC-treated and control NOD mice three weeks after a single dose injection of AAT-MSCs or vehicle and performed scRNAseq analysis (**Fig. 2a**). In the PLN, a total of 8,878 cells from the control group and 8,144 cells from the AAT-MSC group were analyzed. Eight distinct cell clusters were identified based on upregulated gene expression and the expression of key markers specific to each cell population. These include B cells-1 and 2 (identified by *Ms4a*1 expression), CD8^+^ T cells (*Cd8a*^+^), CD4^+^ T cells (*Cd4*^+^), CD4^+^ Treg (*Cd4*^+^ *and FoxP3*^+^), CD4^+^ ISG-EIC (*Cd4*^+^ *and Isg15*^+^), NK cells (*Ccl5*^+^), and macrophages (*Tyrobp*^+^) (**Fig. 2b-d**). Portions of cell populations from control or AAT-MSC groups exhibited differences in each subcell type (**Fig. 2**.**e**).

**Fig. 2.**
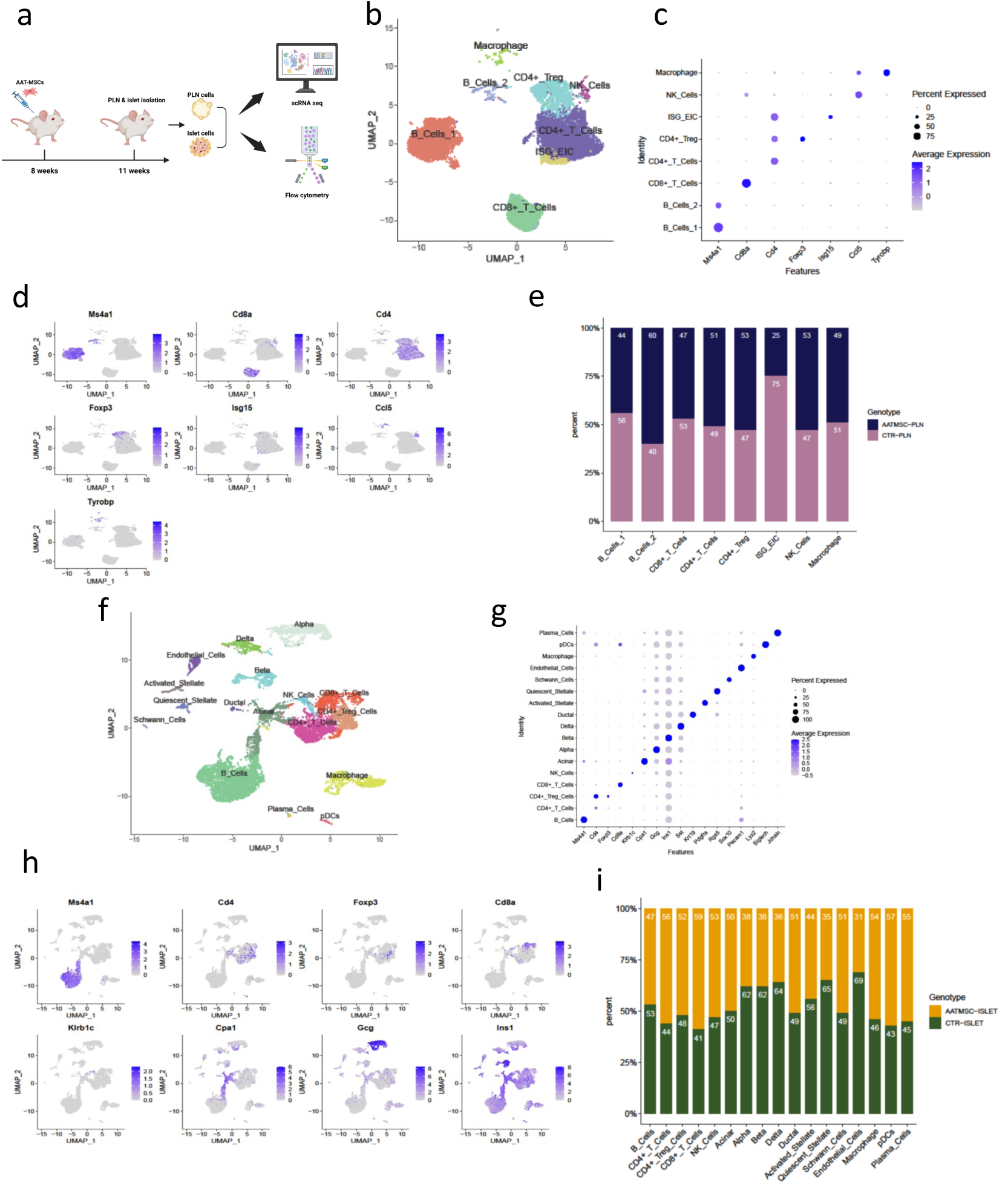
Characterization of different cell types in PLNs and islets from CTR or AAT-treated mice using ScRNAseq analysis. **a**. Schematic of the experimental design for ScRNAseq. ScRNA-seq analysis was performed on PLNs and islets isolated from CTR or AAT-MSC-treated NOD mice at 11 weeks (3 weeks post-treatment, n=3 to 8 per group, samples from individual mice in each were pooled together for ScRNAseq). **b**. t-SNE projection and graph-based clustering of PLN samples from each group. **c**. Gene set enrichment analysis (GSEA) summary of gene signatures for each population in PLN cells. **d**. Expression of canonical cell markers across PLN clusters. **e.** Portion of cells from AAT-MSC or control mice in each cell subtype in PLN cells. **f**. t-SNE projection and graph-based clustering of pooled islet samples from CTR or AAT-MSC-treated NOD mice. **g:** GSEA summary of gene signature for each population in islet cells. **h**. Expression of canonical cell markers across islet cell clusters. **i**. Distribution of cell subtypes in islets from AAT-MSC-treated or CTR mice.

In islets, 6,814 cells were analyzed in the control and 6,422 in the AAT-MSC-treated group. ScRNA-seq identified 17 clusters that highly express markers for immune cells and other pancreatic cell types, including B cells (*Ms4a1*^+^), CD4^+^ T cells (*Cd4*^+^), Treg (*Cd4*^+^*Foxp3*^+^), CD8^+^ T cells (*Cd8a*^+^), nature killer cells (*Klrb1c^+^*), acinar cells (*Cpa1*^+^), alpha (*Gcg*^+^), β cell (*Ins1*^+^*, Ins2*^+^), delta (*Sst*^+^), ductal cell (*Krt19*^+^), and other cell types (**Fig 2. f-h).** Differences in cell portions in both groups were also observed **(Fig. 2i).**

### 3. Characterization of CD4^+^ T cells in the PLN and the islet

Next, we characterized CD4^+^ T cells, key players in the onset and maintenance of T1D, in the PLNs of AAT-MSC-treated and control mice. Using scRNA-seq, we identified six CD4^+^ T cell clusters: CD4 naïve 1 and 2 (*Lef1*^+^), Treg (*Foxp3*^+^), T effector (*Cd69*^+^), IFN-responsive cells (*Stat 1*^+^), and ISG-EIC cells (*Isg15*^+^) in the PLNs (**Fig. 3a-b**). Notably, portions of cell populations from control or AAT-MSC groups exhibited the most dramatic difference in IFN-responsive cells (20% in AAT-MSC-treated vs. 80% in control, **Fig. 3c**).

**Fig. 3.**
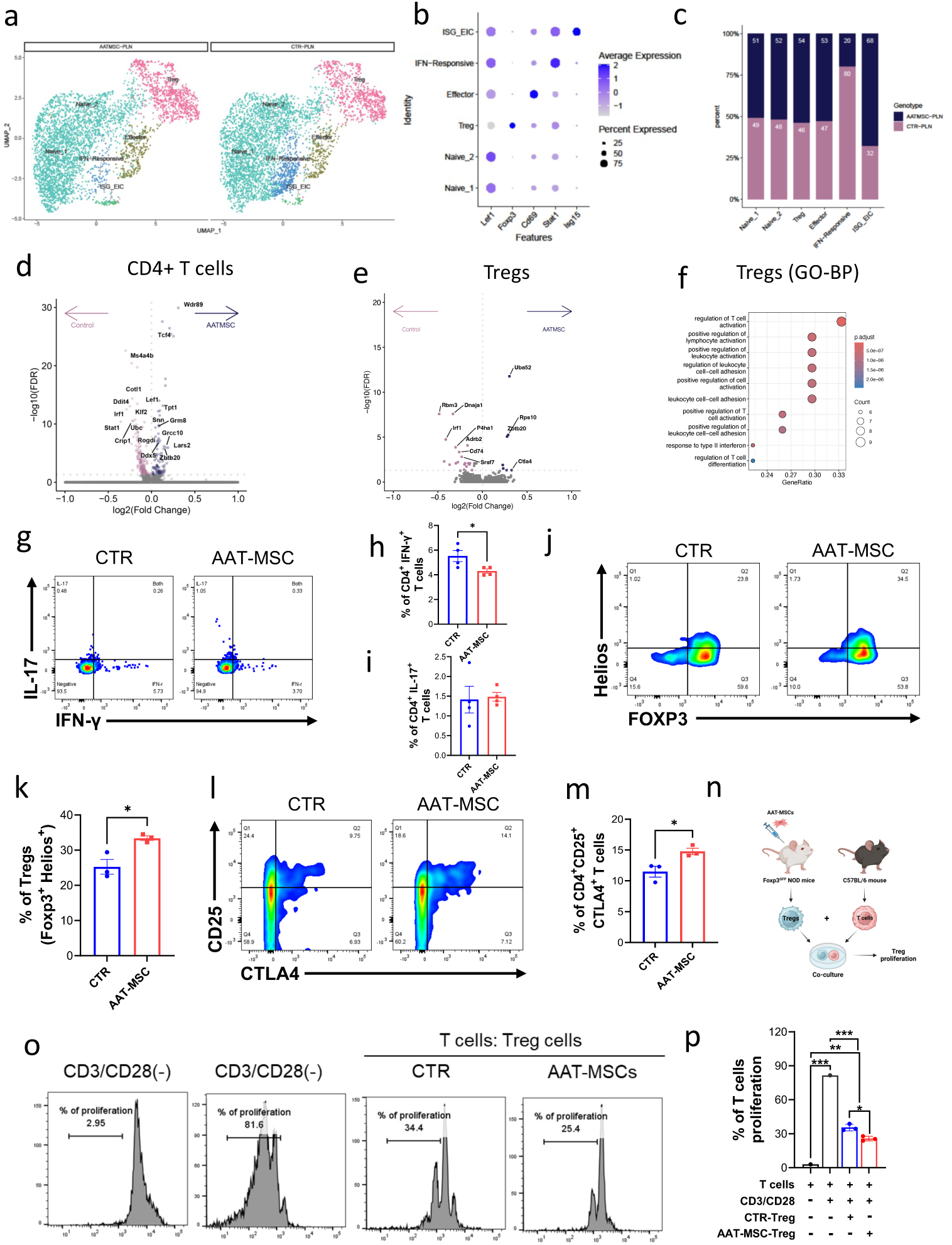
Impact of AAT-MSC treatment on CD4^+^ T cells in the PLN. **a**. t-SNE projection plots depicting CD4^+^ T cell subpopulation in the PLNs of AAT-MSC-treated and CTR mice. **b**. GSEA summary of gene signature associated with CD4^+^ T cell subset. **c**. Distribution of CD4^+^ T subtypes in the PLNs of CTR and AAT-MSC-treated mice. **d &e**. Differentially expressed genes in total CD4^+^ T cells (**d**) and Tregs (**e**) from AAT-MSC-treated or CTR mice. **f.** Gene ontology analysis of the biological process of Tregs in the PLNs. The frequencies of IFN-γ^+^ and IL-17^+^ in CD4^+^ T cells (**g-i**), CD4^+^CD25^+^ Foxp3^+^Helios^+^ (**j&k**), and CD4^+^CD25^+^CTLA4^+^ (**l&m**) were measured and quantified by flow cytometry. N=4 mice per group. *p < 0.05, **p < 0.01. Student’s *t* test. **n.** Schematic diagram of the T cell immunosuppressive assay. Histogram (**o**) and percentage of proliferation (**p**) of mouse T cells cultured with or without anti-CD3/CD28 antibodies in the presence or absence of Tregs from control or AAT-MSC-treated mice (n=6 mice per group). *P < 0.05; **P < 0.01, ***P < 0.001analyzed by Student’s t-test.

The top upregulated genes in total CD4^+^ T cells from the AAT-MSC-group compared to controls were *Wdr89, Rpl35a, Tcf4, RPl13a, Rpl36al*, while the most downregulated genes were *Dnajb1, Ms4a4b*, *Hsp90aa1, Cotl1, and Ddit4, among others* (**Fig. 3d**). Among the upregulated genes, T cell factor 4 (*TCF4)* plays a significant role in regulating immune responses within T cells. Dysregulation of TCF4 has been linked to autoimmune disease, particularly by disrupting the balance between pro-inflammatory and regulatory T cells and limiting the differentiation of pathological effector T cells^38^. RPL13a, a component of the gamma interferon-activated inhibitor of translation (GAIT) complex, is involved in suppressing inflammatory gene translation to control excessive inflammation ^39^. Among the downregulated genes, DNAJB1^+^ T cells play essential roles in developing autoimmune responses. High levels of DNAJB1 expression on T cells can be associated with increased T cell activation and proliferation, potentially enhancing the destructive immune response against pancreatic β cells in T1D^40^. MS4A4B, a membrane adapter protein, is related to antigen response, as naïve T cells transduced with MS4A4B can respond to lower antigen levels ^41^. COTL1 modulates actin dynamics at the T cell immune synapse, which affects T cell spreading and lamellipodial protrusion ^42^. These transcriptional changes suggest that AAT-MSCs regulate the CD4^+^ T cell transcriptome by promoting genes that suppress inflammatory and autoimmune responses while inhibiting genes that enhance immune activation.

A further comparison of the gene expression profiles of the two experimental groups revealed that the top upregulated genes in the IFN-responsive cells are *Rps10, Tcf4, Uba52, Bcl2, and Ptpn20* (**Supplemental Fig. S2**)*. Among these, RPS10* is an IFN-responsive cell component that acts as a negative regulator of the interferon response ^43^. While Bcl2 actively prevents cell death, it can lead to programmed cell death in activated CD4^+^ T cells in the presence of IFN-γ ^44^. *PTPN20* is associated with immune cell infiltration and tumor mutation burden in gastric cancer patients ^45^. Conversely, the downregulated genes were Hsp90aa1 and Hsp90ab1, among others, **Supplemental Fig. S2**).

The frequency of Tregs was comparable between the AAT-MSC and control groups (**Fig. 3c**). Related to gene expression change in the Treg subpopulation, the top upregulated genes in the AAT-MSC group relative to control were *Ctla4*, *Uba52, Rps10, Zbtb20, and Tox* (**Fig. 3e**). UBA52 and Rps10 are ubiquitin-ribosomal fusion proteins that play several essential roles in cellular function ^46^. ZBTB20 plays a crucial role in Tregs, identifying and controlling a subpopulation of Tregs that originate in the thymus and are essential for intestinal homeostasis ^47^. ZBTB20-expressing Tregs have distinct phenotypic and genetic characteristics from non-ZBTB20 Tregs by constitutively expressing IL-10 mRNA and producing high levels of IL-10 upon primary activation. They also express high levels of CD44, TIGIT, GITR, and ICOS compared to non-ZBTB20 Tregs and exert disease-protective effects ^47^. CTLA4 is critical for Treg function ^48,49^. Top downregulated genes in Tregs include *Rbm3, Dnaja1, P4ha1, Irf1, and CD69* (**Fig. 3e**). IRF1 has been shown to negatively regulate Tregs, as IRF1-deficient mice showed increased numbers of Tregs, and the absence of IRF1 leads to enhanced Treg development and function and has implications in autoimmune and inflammatory conditions ^50^. CD69 regulates Treg differentiation and the secretion of IFN-γ, IL-17, and IL-22. It is also a common marker of precursor and mature resident memory T cells (TRMs) localized in peripheral tissues. Functional enrichment analysis of most changed genes in Tregs are involved in the regulation of T cell activation, positive regulation of lymphocyte activation, cell-cell adhesion, and regulation of T cell differentiation (**Fig. 3f** and **Supplemental Fig. S2b&c**). Therefore, treatment with AAT-MSC regulates the expression of key genes that may lead to the improved function of Tregs.

We also characterized CD4^+^ T cells in islets harvested from AAT-MSC-treated and control mice. Three major types of CD4^+^ T cells were observed. These included naïve (*Sell*^+^), regulatory (*Lag3*^+^), and memory (*Itgb1*^+^) CD4^+^ T cells (**Supplemental Fig. S3a&b**). The AAT-MSCs group exhibited more Lag3^+^ Tregs and *Itgb1^+^* memory CD4^+^ T cells than controls (**Supplemental Fig. 3c**). The top upregulated genes in the islet CD4^+^ Treg cells from the AAT-MSC-group were *Rpl38, Rps27, Rps21, Rpl37a,* and *Rps28* (**Supplemental Fig. 3d**). Among them, RPS27 encodes a protein that plays a role in regulating genes associated with the immune response and inflammation ^51^. RPS21 expression is related to immune checkpoints like CTLA4, LAG3, PDCD1, and TIGIT; RPS28 produces damaged-related proteins (DRiPs) on MHC class I molecules, which are critical for T cell recognition ^52^. The most downregulated genes include *Cmss1, Cdk8, Rps6, Ppy,* and *Ins2* (**Supplemental Fig. S3d**). Notably, CDK8 typically promotes the differentiation of anti-inflammatory Tregs while inhibiting Th1 and Th17 differentiation. RPS6 plays a crucial role in T cell development, as its deletion in mouse double-positive thymocytes completely blocks T cell maturation ^53^.

### 4. Treatment with AAT-MSCs suppressed IFN-γ^+^ helper T cells while promoting the immunosuppressive effect of Tregs from the NOD mice

CD4^+^ T helper (Th1) cells are pivotal in immune responses and play a crucial role in T1D pathogenesis by promoting the differentiation, activation, and proliferation of CD8^+^ cytotoxic T cells ^54^. These cells secrete IFN-γ or IL-17, which are associated with Th1 and Th17 responses, respectively ^55,56^. To validate the findings from scRNAseq that indicated a reduction in IFN-γ responsible cells in the AAT-MSC-treated group, we isolated PLNs from NOD mice 3 weeks post-treatment with AAT-MSCs or controls and performed flow cytometry analysis. The results showed that AAT-MSC treatment significantly inhibited the Th1 cell response, as evident by a lower percentage of CD4^+^IFN-γ ^+^ cells in the AAT-MSC-treated PLNs compared to controls (5.52 ± 0.44% in CTR vs. 4.30 ± 0.17% in AAT-MSC, p = 0.04, **Fig 3. g&h**), potentially contributing to the reduced autoimmune response in NOD mice. In contrast, no significant difference was observed in the Th17 cell population between the two groups (1.41 ± 0.33% in CTR vs. 1.48 ± 0.11% in AAT-MSCs, p = 0.85, **Fig 3. g&h**).

Among CD4^+^ T cells, CD25^+^FoxP3^+^ Tregs play a pivotal role in regulating autoimmune responses, a function often compromised in T1D due to a reduced count or impaired functionality of Tregs ^57–60^. To evaluate the impact of AAT-MSC infusion on Treg number and function, we quantified the percentages of CD4^+^CD25^+^FoxP3^+^ Tregs in the PLNs from both the AAT-MSC-treated and control NOD mice, 3 weeks post-treatment. As shown in the ScRNAseq analysis, we observed no significant difference in the overall number of Tregs in the PLNs of the two groups (6.94 ± 0.93% in CTR vs. 7.02 ± 0.48% in AAT-MSC, p = 0.88, **Supplemental Fig. S2d&e**). However, AAT-MSC-treated mice exhibited a significant increase in Foxp3^+^Helios^+^ (25.27 ± 2.09% in CTR vs. 33.43 ± 0.63% in AAT-MSC, p = 0.02, **Fig 3. j&k**) and CD4^+^CD25^+^CTLA4^+^ cells (30.50 ± 1.60% in CTR vs. 49.17 ± 6.42% in AAT-MSC, p = 0.047, **Fig 3. L&m)**. These two Treg subpopulations are associated with enhanced immunosuppressive function ^48,49^.

To determine whether AAT-MSCs enhanced the immunosuppressive function of Tregs, Tregs were isolated from the PLNs of female Foxp3^EGFP^ NOD mice treated with AAT-MSCs or controls and co-cultured with T cells from C57BL/6 mice using a standard T cell proliferation assay (**Fig. 3n**). Upon stimulation with anti-CD3 and anti-CD28 antibodies, T cells showed a rapid proliferation rate of 85.50 ± 2.38%. When co-cultured with Tregs from control NOD mice, T cell proliferation was reduced to 35.50 ± 2.38%. In contrast, T cells co-cultured with AAT-MSC-derived Tregs exhibited a further reduction in proliferation to 25.70 ± 1.65% (n = 6 mice per group, p < 0.05 vs control Tregs, p < 0.001 vs. T cell with CD3/CD28, ANOVA, **Fig 3. o&p**). These results suggest that treatment with AAT-MSCs enhances the immunosuppressive function of Tregs, likely due to the increased expression of Helios and CTLA4 in these cells.

### 5. Co-culture with AAT-MSCs increases mouse and human Treg numbers in vitro

We next investigated whether co-culturing immune cells with AAT-MSCs could increase Treg numbers *in vitro,* both in mouse and human samples. First, we co-cultured mouse CD4^+^ T cells with AAT-MSCs, with or without anti-CD3 and CD-28 antibodies, for three days. After this period, we measured the proportion of CD4^+^CD25^+^Foxp3^+^ Tregs by flow cytometry. Our data demonstrated that co-culturing with AAT-MSCs not only reduced the total number of CD4^+^ T cells but also significantly increased Treg numbers and percentages, regardless of CD3/CD28 stimulation (**Fig. 4. a-c**). This increase in Tregs was likely driven by enhanced Treg proliferation, as indicated by cell proliferation analysis (**Fig. 4d**). Additionally, we observed significant increases in Foxp3^+^Ctla4^+^ and Foxp3^+^Helio^+^ subpopulations of Tregs in cells treated with AAT-MSCs **(Fig. 4&f)**, further confirming that AAT-MSCs not only promote Treg proliferation but also the expressions of immunosuppressive markers including CTLA4 and Helios, as observed *in vivo*. Next, we evaluated whether this effect could be replicated in human cells. We co-cultured peripheral blood mononuclear cells (PBMCs) from healthy donors with AAT-MSCs for three days and measured the percentages of the CD4^+^CD25^+^CD127^low^ Treg population (**Fig. 4g**). PBMCs cultured alone exhibited a Treg population of 4.09 ± 0.58%. In contrast, PBMCs co-culture with AAT-MSCs at a 1:10 or 1:5 ratio showed significant increases in the Treg population, up to 6.21 ± 0.14% and 6.72 ± 0.22%, respectively (**Fig 4. h&i**). These results prove that co-culture with AAT-MSCs increases mouse and human Tregs *in vitro*.

**Fig. 4.**
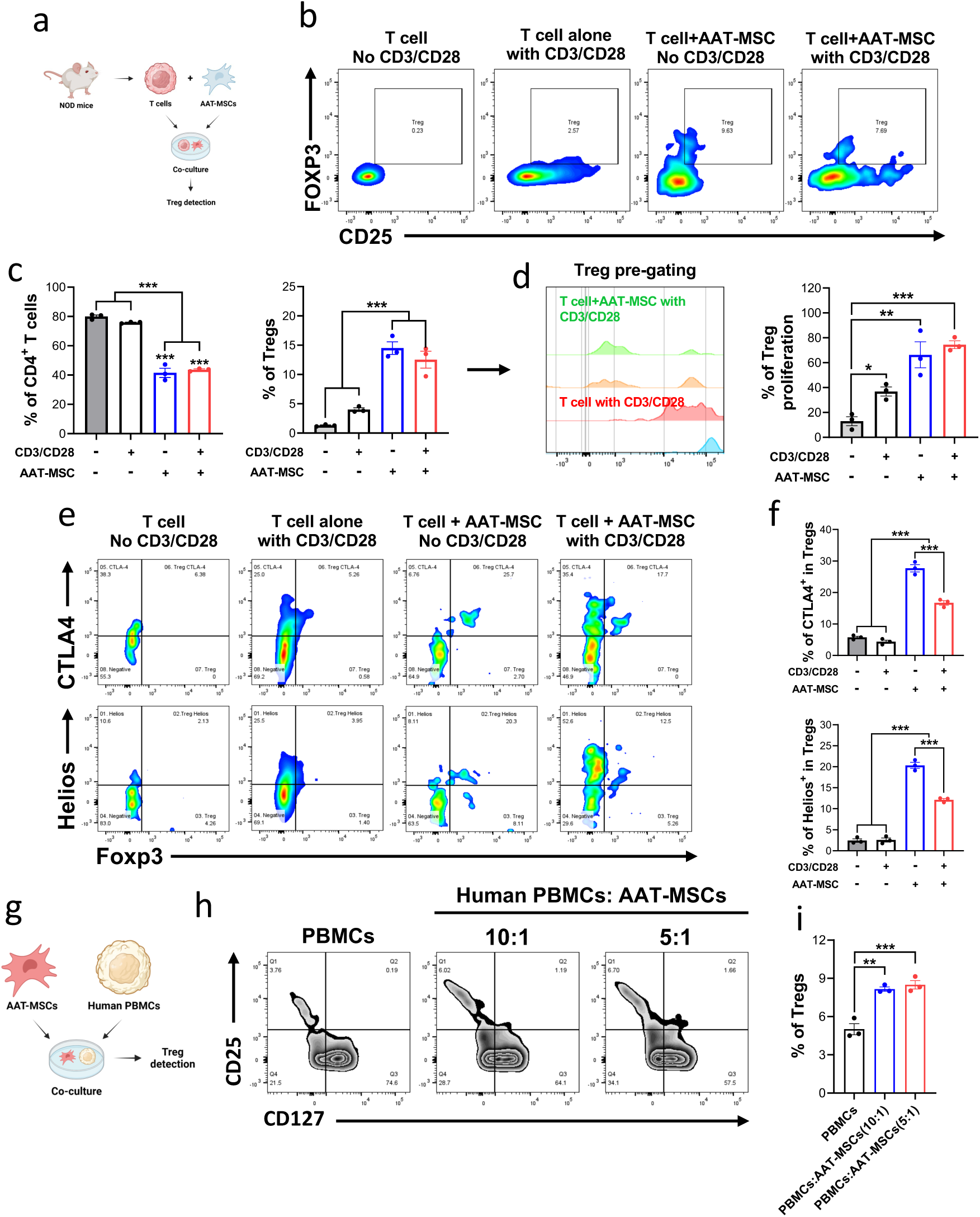
Co-culture with AAT-MSCs increases Tregs among T cells from NOD mice and human PBMCs. **a.** Schematic of AAT-MSCs and T cell co-culture. **b&c**: Scatter dot plots **(b**) and frequency (**c**) of mouse CD4^+^CD25^+^Foxp3^+^ Tregs, as well as CTLA4^+^ and Helio^+^ Treg (**e and f**) in cultures with or without anti-CD3/CD28 antibodies and/or AAT-MSCs. (**d**). Histogram shows Treg cell proliferation and percentage of proliferation. **g**. Schematic of human Tregs induced by co-culture of PBMCs with AAT-MSCs. **h, i**. Analysis of CD4^+^CD25^+^CD127^lo^ Treg population in human PBMCs co-cultured with AAT-MSCs. Data are presented as mean ± SEM of at least three independent experiments, and the scatter dot plot displays individual data point. *p < 0.05, **p < 0.01, ANOVA test.

### 6. Treatment with AAT-MSCs favors exhausted CD8^+^ T cell phenotype in the islets

CD8^+^ T cells are the primary immune cells that infiltrate the islets of NOD mice and T1D patients and play a significant role in the pathogenesis of T1D by directly killing β cells ^61–63^. To investigate the impact of AAT-MSC treatment on CD8^+^ T cell profiles, we performed ScRNAseq analysis in islets of AAT-MSC-treated and control NOD mice at 3 weeks post-treatment. We identified three major sub-clusters of CD8^+^ T cells in the islets: proliferative (*Mki67^+^*), memory (*CD28^+^*), and exhausted (*Tox^+^*) CD8^+^ T cells. The islets of AAT-MSC-treated mice exhibited higher proportions of all three subpopulations (**Fig. 5 a-c**).

**Fig. 5.**
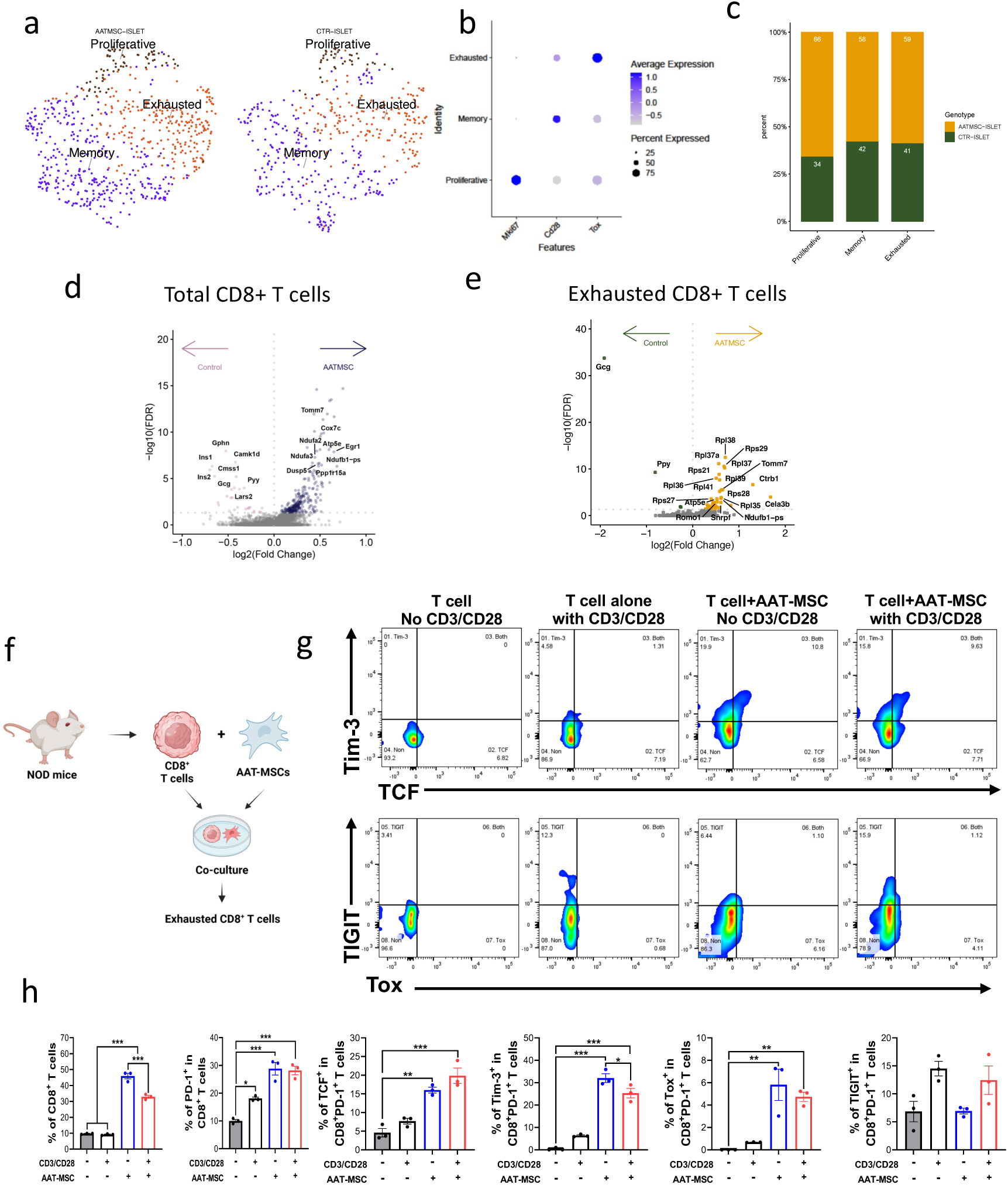
AAT-MSCs increase exhausted CD8^+^ T cells in vivo and in vitro. **a**. t-SNE projection plots of CD8^+^ T cells in the islet from the mice treated with AAT-MSC or CTR. **b**. GSEA summary of gene signature in subtypes of CD8^+^ T cells. **c**. Portions of the subtype population of CD8^+^ T cells in the islet from AAT-MSC-treated or CTR mice. **D& e** Differentially expressed genes in total and exhausaed (Tox^+^) CD8^+^ T cells. **f.** Schematic measuring mouse exhausted CD8^+^ T cell induced by AAT-MSC co-culture *in vitro*. **g, h**. Scatter dot plot (**g**) and quantification (**h**) of exhausted CD8^+^ T cells induced by co-culture with AAT-MSCs. Isolated T cells were co-cultured with AAT-MSCs for 5 days, followed by antibody staining to detect exhausted CD8^+^ T cells. CD8^+^ PD-1^+^ cells were pre-gating, and representative flow cytometry data show the expression of exhausted CD8^+^ T cells marker, including TCF, Tim-3, Tox, and TIGIT, in the presence or absence of AAT-MSCs. Data are presented as mean ± SEM from at least three independent experiments, with individual data points shown in the scatter dot plot. *p < 0.05, **p < 0.01. One-way ANOVA with post-hoc correction.

The top upregulated genes in total CD8^+^ T cells from the AAT-MSC islets compared to control islets include *Tomm7, Cox7c, Egr1, and Ndula2*, while downregulated genes include *Camk1d, gphn, and Cmss1, among others* (**Fig. 5d**). Tomm7 or “let-7” is associated with CD8^+^ T cell activation, as decrease of let-7 expression in activated T cells enhances clonal expansion and the acquisition of effector function ^64^. Among the downregulated genes, CAMK1D is expressed in CD8^+^-activated T cells, not in resting cells. Its activity increases upon T cell activation, where it plays a key role in CD8^+^ T cell proliferation, cytotoxic activity, and responsiveness to stimulation ^65^.

Next, we focused on analyzing the exhausted CD8^+^ T cell subpopulation because exhausted CD8^+^ T cells have been correlated with slow progression of and response to treatment in immunotherapy trials for T1D, such as anti-CD3 therapy (teplizumab) ^1,66,67,68^. The upregulated genes in CD8^+^ exhausted T cells from the AAT-MSC group include *Ctrb1, Rpl35, Rpl37a,* and *Rpl38,* and the top downregulated genes include *Camk1d* and *Gcg (****Fig. 5e***). Our data suggest AAT-MSCs promote CD8^+^ T cells resting with low cytotoxic capability, reflecting their exhausted feature.

In addition, the characterization of CD8^+^ T cells in the PLN showed there were four major CD8^+^ T cell populations: memory (*Il7r^+^),* cytotoxic (*Nkg7^+^*), exhausted (*Tcf7^+^ or Tox^+^*) **(Supplemental Fig. S4 a&b)**. There were fewer *Il7r^+^* memory CD8^+^ T cells in the AAT-MSCs compared to controls, while the numbers of the other three types of CD8^+^ T cells are comparable **(Supplemental Fig. S4c).** IL-7R^+^CD8^+^ T cells play a role in the autoimmune destruction of pancreatic β cells in T1D. These cells are involved in the disease’s progression, as they can become autoreactive and target the insulin-producing β cells, contributing to the development of the condition ^69^. AAT-MSC treatment likely suppresses CD8^+^memory T cells, thus protecting β cell death.

We compared gene expression in the Tox^+^ exhausted CD8*^+^* T cell subpopulation **(Supplemental Fig. S4d)**. Specifically, the expression of Ms4a4b and lef1, both markers for exhausted CD8^+^ T cells, was upregulated. In contrast, Stat1 and Klf2, critical factors for CD8^+^ T cell proliferation and function, were downregulated in exhausted CD8^+^ T cells from AAT-MSC-treated cells. These gene expression changes suggest that AAT-MSCs promote an exhausted CD8^+^ phenotype.

We next deciphered the impact of AAT-MSCs on exhausted CD8^+^ T cells *in vitro*. Mouse splenocytes were co-cultured *ex vivo* with AAT-MSCs in the presence or absence of anti-CD3 and anti-CD28 antibodies, and the frequency of exhausted CD8^+^ T cells was assessed **(Fig. 5f)**. Co-culturing splenocytes with AAT-MSCs increased the frequency of both total CD8^+^ T cells and PD-1^+^ exhausted CD8^+^ T cells **(Fig. 5g&h**). The frequency of progenitor-exhausted CD8^+^ T cells (characterized by the expression of PD-1, TCF, and Tim3) and terminal exhausted CD8^+^ T cells (identified by expression of PD-1, Tox, and/or TIGIT) were significantly higher in the AAT-MSCs-co-culture group compared to those from controls (**Fig. 5h**). These findings suggest that AAT-MSC promotes the transition of T cells into a CD8^+^ T cell exhaustion phenotype.

### 7. Tregs from AAT-MSC-treated islets established cell-to-cell interactions with other cells in islets, likely via the enhanced IL-2 signaling

Utilizing the CellChat application, we analyzed cell-cell communications in PLN and islet cells using ScRNAseq data. Cells in control PLNs exhibited fewer interactions (75 interactions) than those from AAT-MSC PLNs (104 interactions, **Fig. 6a&b, supplemental Fig. S5a**). Among the incoming signaling pathways in PLN cells, TGF-β, a key Treg differentiation factor, exhibits the highest relative strength in Tregs from AAT-MSC-treated mice compared to controls, but not much difference in the outgoing pathways (**Supplemental Fig. S5b-d**).

**Fig. 6.**
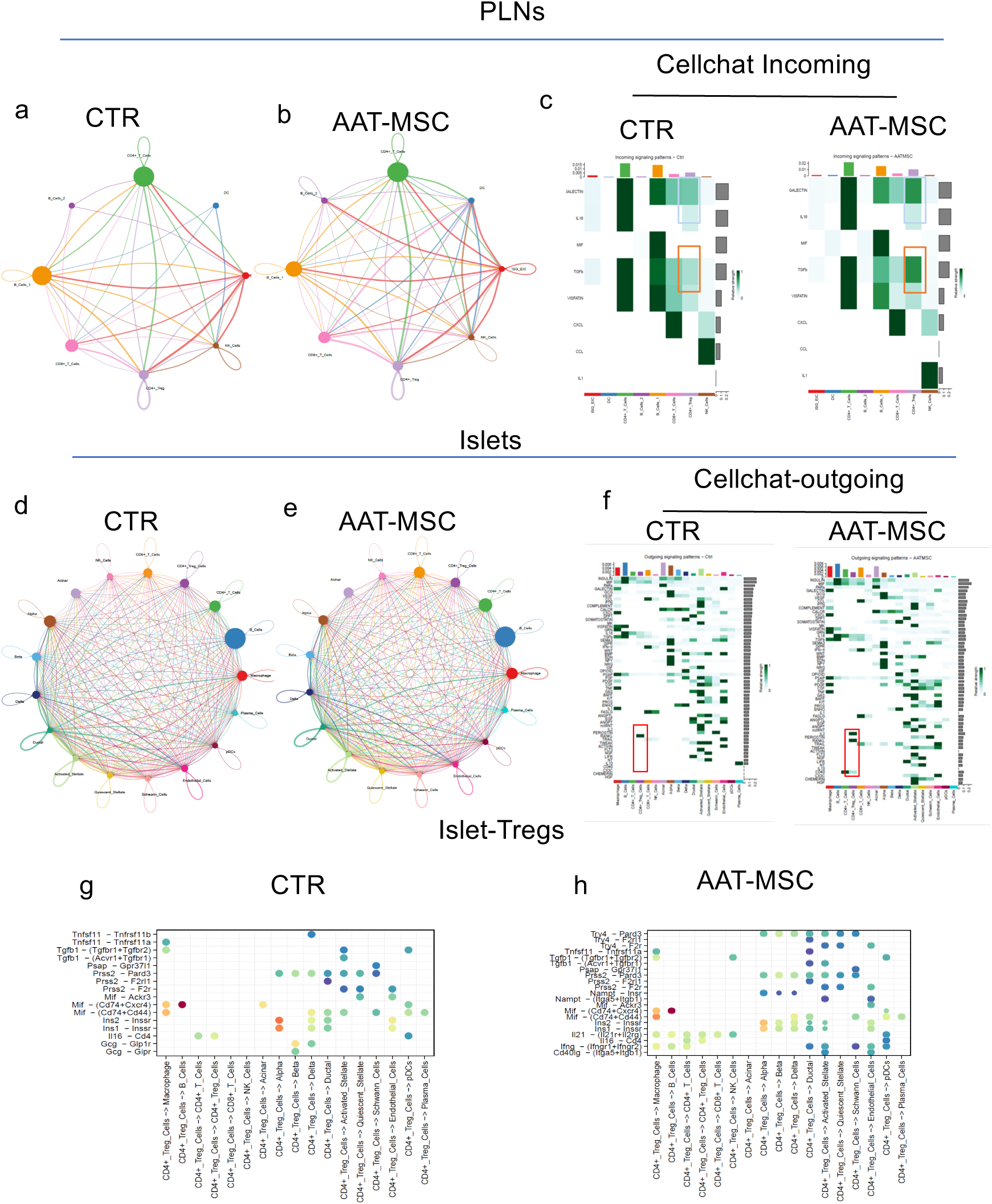
Enhanced cell-cell communications in PLNs and islets from AAT-MSC-treated mice compared to controls as analyzed by Cellchat. **a.** Netvisual aggregate of cells from PLNs of CTR (**a**) or AAT-MSC (**b**) mice. **c.** Incoming signals in immune cell subtypes from PLNs of CTR or AAT-MSC mice. d. Netvisual aggregate of cells from islets of CTR (**d**) or AAT-MSC (**e**) mice. f. Outgoing signals in immune cell subtypes from islets of CTR or AAT-MSC mice. g: Dot plots showing ligand-receptor prediction analysis between CD4^+^ Tregs and other immune cells in islets from CTR or AAT-MSC-treated mice.

In the islets, control islet cells showed 2,211 interactions, while AAT-MSC-treated islet cells have 2,593 interactions (**Fig. 6d &e and supplemental Fig. S5e**). Notably, AAT-MSC islets showed enhanced IL-2 outgoing signaling (**Fig. 6f**). Furthermore, compared to controls, Tregs from the AAT-MSCs treated islets are highly interactive with the microenvironment and establish a more complex cell-to-cell communication network with other cell types via multiple pathways as shown by the increased outgoing signaling (**Fig. 6g&h**) as well as ligand-receptor prediction analysis by Cellchat (**Supplemental Fig S5f**). Overall, our results reveal that treatment with AAT-MSCs enhances cell communication, particularly from Tregs to other cell types in the PLN and islet.

### 8. Tregs educated by AAT-MSCs promote the conversion of effector CD8^+^T cells into an exhausted phenotype

We investigated whether AAT-MSC-educated Tregs can directly promote the generation of an exhausted CD8^+^ T cell phenotype based on their enhanced communication with other cells. We separated CD4^+^GFP^+^ Tregs and CD4^+^GFP^-^ non-Tregs from spleen and PLN cells from the Foxp3^EGFP^ NOD mice and co-cultured them with or without AAT-MSCs for 72 hours. Subsequently, all four groups of cells were co-cultured with mouse CD8^+^ T cells for an additional 72 hours, and the percentages of exhausted CD8^+^ T cells were measured by flow cytometry. In some experiments, these cells were co-cultured with freshly isolated mouse islets to assess their impact on islet cell death (**Fig. 7a**).

**Fig. 7.**
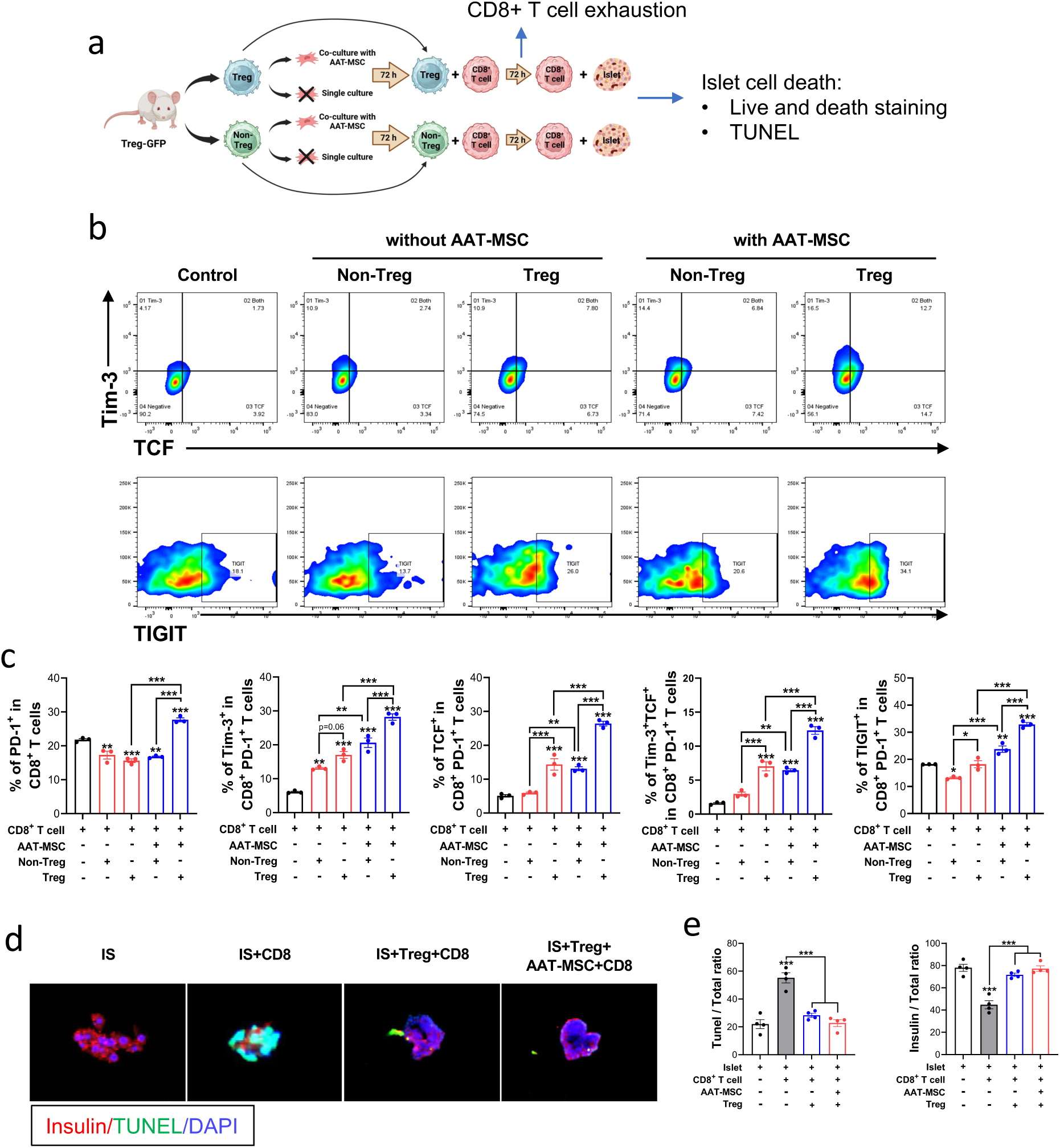
AAT-MSC-educated Tregs induce exhausted CD8^+^ T cell phenotypes that promote human islet survival ex vivo. **a.** Schematic illustration of Treg education with AAT-MSCs, its impact on CD8^+^ T cell exhaustion and islet suvival. (**b and c**) Scatter plots (**b**) and percentages (**c**) of exhausted CD8^+^ T cells expressing PD-1^+^, Tim-3, TCF, and TIGIT in the presence or absence of AAT-MSC co-culture for 72 hours were analyzed by flow cytometry. **d-e**: Microscope (**d**) images and quantification (**e**) of human islet cell death (measured by TUNEL^+^/total islet cells) and insulin^+^ cells/total islet cells in the presence or absence of CD8^+^ T cell and/or AAT-MSCs co-culture, quantified by TUNEL assay and insulin staining. Red: insulin^+^ cells, green: TUNEL^+^ cells, blue: DAPI. The bar graph represents the mean ± standard error mean (SEM), and the scatter dot plot represents an individual data point. * p< 0.05; **, p < 0.01, ***, P < 0.001. NS; not significant, ANOVA test.

Our data revealed that expression of the exhaustion marker, PD-1, Tim3, ICF, TIGIT were elevated at different extents in CD8^+^ T cells in the groups co-cultured with GFP^+^Tregs or AAT-MSC-educated Tregs, with the latter showing more pronounced effects. In contrast, GFP^-^ non-Tregs alone had little, if any, effects on CD8^+^ T cell exhaustion. However, non-Tregs educated by AAT-MSCs showed an increase in exhausted CD8^+^ T cells compared to non-Tregs cultured without AAT-MSCs (**Fig. 7b&c**). These findings suggest that CD4^+^Tregs, but not non-Tregs, can directly promote CD8^+^ T cell exhaustion, with AAT-MSC education enhancing this effect in both Tregs and non-Tregs, particularly in Tregs. Furthermore, TUNEL staining of islets (**Fig. 7 d&e**) and a live and dead cell analysis (**Supplemental Fig. S6**) confirmed a positive correlation between islet cell survival and the percentage of exhausted CD8^+^ T cells: higher levels of islet cell death were observed when islets were co-cultured with CD8^+^ T cells or non-Treg cells, whereas islet survival was significantly better in islets co-cultured with AAT-MSC-educated Tregs. These findings suggest that AAT-MSCs primarily protect islet survival through Tregs and their abilities to convert CD8^+^ Tcells into an exhausted phenotype that allows islet cell survival.

### 9. AAT-MSC infusion results in the reversal of hyperglycemia via increasing the number and/or function of Tregs and suppressing insulitis and T cell activation

We expanded our therapeutic approach to tackle a more challenging T1D mouse model, where we administered a single dose of AAT-MSCs to female NOD mice exhibiting newly onset T1D (with two consecutive blood glucose >250mg/dl). Untreated NOD mice (n=11) displayed a drastic increase in blood glucose levels over time. In contrast, mice treated with AAT-MSCs (n=18) demonstrated considerably lower average blood glucose levels and a significantly smaller AUC of blood glucose levels (p<0.0001, vs. control, **Supplemental Fig. S7a&b**). Moreover, they experienced a significantly prolonged duration of normoglycemia post-treatment (**Supplemental Fig. S7c**). In addition, mice treated with AAT-MSCs exhibited an increased number of CD4^+^CD25^+^FoxP3^+^ Tregs cells and higher percentages of CD4^+^CD25^+^Ctla^+^ Tregs (**Supplemental Fig. S7, d-g).** AAT-MSC infusion also led to a significant reduction in insulitis. Control NOD mice showed that 28.2 ± 3.6% of islets had <5% immune cell infiltration, and 50.6 ± 4.3 % had over 75% infiltration. In contrast, AAT-MSC treatment resulted in 51.1 ± 4.3 % of islets showing <5% immune cell infiltration, while 26.2 ± 4.2% exhibited infiltration of 75% or more (**Supplemental Fig. 7S, h-i**).

Additionally, PLN cells isolated from mice receiving AAT-MSCs showed significantly lower productions of IFN-γ, IL-6, TNF-α, and IL-1β and an overall higher IL-10: IFN-γ concentration ratio after *ex vivo* activation by anti-CD3 antibody compared to cells from non-treated control NOD mice (**Supplemental Fig. S7, j-n**). This data suggests that AAT-MSCs can lead to a reversal of hyperglycemia in mice with newly onset T1D, likely via promoting Tregs cell survival and function and reducing immune cell infiltration and activation within the islets.

## DISCUSSION

The therapeutic effects of naïve or AAT-overexpressing MSCs in T1D and GvHD have been described in mouse models and clinical trials ^21,34,36,70^. However, the mechanisms of how these cells suppressed autoimmunity and re-established homeostasis *in vivo* to halt destruction to pancreatic β cells are incompletely explored. In this study, we demonstrated that a single infusion of AAT-MSCs achieved a profound therapeutic effect via modulating subtypes of immune cells critical for T1D. AAT-MSC infusion led to an increase in the immunosuppressive function of Tregs by enhancing Helios and CTLA4 expression, suppressing IFN-responsive (*Stat1^+^*) cells. Meanwhile, AAT-MSC infusion is correlated with more exhausted CD8^+^ T cells and contributes to β cell survival. *In vitro*, co-culturing splenocytes or PBMCs promoted T cell Treg generation and led to CD8^+^T cell exhaustion. Importantly, these effects translated in the NOD mice with newly onset T1D, in which AAT-MSC therapy reversed diabetes in some mice. Our comprehensive data demonstrated that AAT-MSCs can be used for T1D therapy.

AAT-MSCs enhanced the immunosuppressive effects of Tregs. Treg insufficiency and dysfunction were observed in T1D patients and NOD mice, and increased Tregs are associated with delay in T1D progression in children ^71^. There are persistent efforts to stimulate Tregs and suppress T effector cells to protect β cells, and direct infusion of Tregs has been shown to benefit T1D therapy ^72^. Tregs use several mechanisms for immune regulation, including the production of the anti-inflammatory cytokine IL-10, depletion of IL-2 needed for the proliferation of effector T cells, and transmission of inhibitory signals to both APCs and T cells via CTLA-4 ^73^. We found that AAT-MSC infusion enhances the immunosuppressive function of Tregs by promoting the expressions of Helios and CTLA4, two critical transcriptional factors for Treg function. Helios expression can ensure stable expression of a suppressive and anergic phenotype in the face of intense inflammatory responses ^74^. This aligns with research conducted by other groups, which demonstrated that MSCs can stimulate the growth of Tregs by releasing certain factors and reinstate the disrupted Th17/Treg balance in autoimmune disease ^75,76^. Furthermore, AAT-MSCs, either through cell-cell interaction, secretion of IL-10, or AAT, increase Treg populations and/or function, both *in vivo* in the NOD mice and *in vitro* when co-cultured with mouse splenocytes or human PBMCs. This is likely through promoting the expression of key genes related to Treg stability and/or function, conversion of T effector cells into Tregs, or protection of Treg from cell death. This function could provide insights into the increased immunosuppressive function of Tregs observed *in vivo*, potentially contributing to the therapeutic effects of AAT-MSCs in modulating immune responses.

Additionally, treatment with AAT-MSCs suppressed IFN-responsive T cells. Previous studies have shown that human umbilical cord-derived MSCs (hUC-MSCs) inhibit STAT1/3 signaling in T cells by secreting chitinase-3-like protein 1 (*CHI3L1*) and upregulating peroxisome proliferator-activated receptor δ (PPARδ). MSCs interfere with the phosphorylation of pSTAT1 and its binding to IFN-stimulated response elements, thereby controlling the expression of interferon-stimulated genes ^77^. This mechanism is consistent with our finding that AAT-MSCs reduce Th1 cells in the PLN. Our study proves that much of the therapeutic effects in T1D are due to AAT-MSCs’ inhibitory effect on *Stat1^+^* IFN-responsive cells. The emergence of CD4^+^ T cell subsets detrimental to pancreatic β cell survival plays a crucial role in T1D. The canonical Th1 cytokine, IFN-γ, contributes to Th1 cell development, as evidenced by studies showing that mice deficient in IFN-γ (IFN-γ^−/−^) exhibit aberrant Th2 cell development when faced with pathogens that typically induce Th1 responses ^78^. IFN-γ influences Th1 development by regulating IL-12Rβ2 chain expression and promoting IL-12 secretion from macrophages directly or through activated NK cells ^79^. In addition, IFN-γ produced by Th1 cells can suppress the growth of Th2 cells ^80^. In our study, AAT-MSC therapy suppressed the IFN-responsive Th1 cells in the PLN of mice: there was a significant decrease in IFN-responsive cells within the PLNs treated with AAT-MSCs compared to controls. This subpopulation of cells is characterized by elevated expression levels of the transcription factors *Stat 1* and *Lef1* alongside the membrane-bound type II C-lectin receptor CD69. Stat1 is a transcriptional factor that mediates signaling downstream of IFN-α, IFN-β, and IFN-γ. CD69 is known for its rapid appearance on lymphocyte surfaces following activation and is an early marker of lymphocyte activation ^81^; it associates with a predisposition to autoimmune conditions and affects Th/Treg balance and the suppressive activity of Tregs ^82^.

We did not observe an effect of AAT-MSCs on circulating cytokine levels in the T1D prevention model, but the suppressive effects were more dramatic in the T1D treatment model. This is probably because the levels of Th1-related IFN-γ and Th2-related IL-4 were increased. Similarly, we did not observe Treg number changes in the prevention model; but Treg numbers were increased in the treatment model. Therefore, both number and function appear necessary since enhanced Treg functions were also observed in both models, highlighting the potential effect of AAT-MSCs in treating human T1D.

Another novel finding is that AAT-MSC treatment favors a CD8^+^ exhausted T cell phenotype characterized by increased PD-1, TCF, Tim3, Tox, and TIGIT expression in the CD8^+^ T cells within islets as identified by ScRNAseq and confirmed *in vitro*. In humans, the exhausted cells are hyperproliferative and share expanded rearranged T cell receptor junctions and expressed exhaustion-associated markers, including TIGIT and KLRG1 ^69^. In both the NOD mice and humans, CD8^+^ T cells are a major component of the immune infiltration in the islets. The islet antigen reactive CD8^+^ T cells can also be reproducibly detected in the blood and pancreas of patients with T1D ^69^. In recent years, exhausted CD8^+^ T cell signatures have been linked with slower progress of T1D after diagnosis and clinical response to immunotherapy ^67^. For example, two distinct CD8^+^ T cell signatures exist at different stages of T1D: a proinflammatory signature in children with newly diagnosed T1D and a co-inhibitory signature in autoantibody-positive children who later progressed to T1D, which suggests that CD8^+^ T cells signatures could potentially be used as biomarkers for evaluating T1D progress ^69^. Thus, our data shows for the first time that AAT-MSCs promote the exhaustion of CD8^+^ T cells that might have contributed to islet survival in the NOD mice.

Our results also demonstrated novel insights into the role of AAT-MSC-educated Tregs in promoting the exhaustion of CD8^+^ T cells and their subsequent protective effect on islet survival. The data that Tregs, particularly those educated by AAT-MSCs, upregulated exhaustion markers CD8^+^ T cells suggest that Tregs, but not non-Tregs, can directly induce CD8^+^ T cell exhaustion. Additionally, our data showed a correlation between the exhaustion of CD8^+^ T cells and enhanced islet cell survival in the in vitro cell culture system. Therefore, AAT-MSCs confer a protective effect on islet survival mainly through Tregs, which promote CD8^+^ T cell exhaustion and reduce islet cell death. These data suggest the potential therapeutic applications of AAT-MSCs in autoimmune diseases and transplantation. The *in vitro* studies suggest that AAT-MSCs can enhance the numbers of Tregs derived from PBMCs, possibly through multiple mechanisms: direct conversion of T effector cells into Tregs and protection of Tregs from cell death. These functions could provide insight into the increased immunosuppressive function of Tregs observed *in vivo*, potentially contributing to the therapeutic effects of AAT-MSCs in modulating immune responses. Taken together, under autoimmune diabetes conditions, the function of Tregs decreases while the activity of T effector cells increases, leading to islet cell death. However, when AAT-MSCs are given, they boost Treg function and reduce effector T cell activity. Tregs enhanced by AAT-MSCs can also transform CD8^+^ T cells from an effector phenotype to an exhausted phenotype. These exhausted CD8^+^ T cells may interact with other impaired effector T cells, favoring the survival of pancreatic islet cells.

Our studies have limitations. We pooled PLN or islet cells from multiple mice for scRNAseq, as RNA sample pooling strategies can optimize both the cost of data generation and statistical power for differential gene expression analysis ^83^. PLNs and islets were analyzed 3 weeks post cell infusion time point because we anticipated that this would be the timeframe for observing the impact of MSCs, making it an optimal time to compare the cellular landscape in the PLNs and islet between control and treated NOD mice. However, additional times may show more dramatic effects. In this study, we only focused on T cells, although other cell types may also contribute to the outcome of AAT-MSC protection. As a T cell-driven autoimmune disease that targets β cells, an essential aspect of treatment aims to dampen autoimmune attacks, especially among individuals in the early stages of T1D. Both MSC and AAT showed profound protection in autoimmune diseases. Therefore, the effects observed may be a combination of both MSCs and AAT. AAT-MSCs may prolong survival, enhance the function of MSC, and induce sustained suppression of autoimmunity and β cell protection. MSC and AAT target different problems in T1D, and AAT-engineered MSCs show dramatically improved functionality compared to control MSCs with consistent secretion of AAT at a high level. Additional studies are needed in some aspects of our work to determine the relative contribution of the AAT gene edits compared to AAT or MSC treatment alone.

In summary, through comprehensive *in vitro* and *in vivo* analyses, we identified major T cell subpopulations and intracellular signaling events related to the effects of AAT-MSCs on diabetes remission or progression. AAT-MSCs achieve therapeutic effects in T1D by suppressing autoreactive T cell infiltration and activation, enhancing Treg function, and promoting CD8^+^ T cell exhaustion. These mechanisms collectively contribute to the protection against T1D progression.

## MATERIALS AND METHODS

### Preparation of AAT-MSCs

Human bone marrow-derived MSCs were isolated from human bone marrows and infected with lentivirus to overexpress AAT, as described previously ^34^. AAT-MSCs were incubated at 37°C in 5% CO2 in *α*-MEM (Gibco, cat. A10490-01) supplemented with 10% fetal bovine serum (FBS; Gibco, cat. A52568-01), 2 mM L-glutamine (Gibco, cat. 25030-081), 100 U/mL penicillin (Gibco), and 100 mg/mL streptomycin (Gibco). Once the cultures reached 90% confluency, cells were subcultured or stored in liquid nitrogen. Cells at passage 6∼8 were used in this study.

### Mice and AAT-MSCs infusion

Female NOD mice were obtained from the Jackson Laboratory (Bar Harbor, ME). Two treatment models were employed in this study. The first was the diabetes prevention model, in which mice aged eight weeks received a single intravenous infusion of AAT-MSCs with a dose of 1x10^6^ cells per mouse. The second model involved newly diagnosed T1D mice, where NOD mice with two consecutive non-fasting blood glucose readings > 250 mg/dl were administered a single dose of AAT-MSCs at 1x10^6^ cells per mouse. Age-matched female NODs were used as controls. Non-fasting blood glucose levels of mice were measured by tail vein prick using the Freestyle Lite glucometer (Abbott Inc) twice weekly.

All mouse studies were approved by the Institutional Animal Care and Use Committee at the Medical University of South Carolina.

### Intravenous glucose tolerance test (IVGTT)

Mice were fasted for 4 hours before receiving a glucose injection at 1g/kg body weight via the tail vein. Blood glucose levels were measured at baseline, 2, 15, and 30 minutes after the glucose infusion. A small aliquot of blood was collected at each time point to obtain serum for C-peptide measurement using a mouse C-peptide ELISA kit (APLCO) according to the manufacturer’s recommendation.

### Hematoxylin and eosin (H&E) staining of pancreas

The whole pancreas was isolated, fixed, and embedded in paraffin. At least three sections spanning every 100 μm of the pancreas were selected from serially sectioned slides and stained for H&E using a standard protocol. Insulitis scores were graded as follows: grade 0, a normal islet with <5% of immune cell infiltration; grade 1, 5-25% of the islet were infiltrated by immune cells; grade 2, 25–50% of the islet were infiltrated; grade 3, 25-50% of the islet were infiltrated; grade 4, >75% of islet were infiltrated. Each islet was evaluated by at least three people independently. Data were pooled from sections obtained from different mice in each group.

### RNA isolation and RT-PCR analysis

Total RNA was isolated from cells using Trizol reagent (Invitrogen), and then reverse-transcribed by M-MLV reverse transcriptase with oligo (dT) 18 primers (Thermo Fisher). Real-time PCR was performed using SYBR Green I (Thermo Fisher) on a CFX96 Real-Time PCR Detection System (Bio-Rad). The thermal profile for qPCR was 95 °C for 10 min, followed by 40 cycles of 95 °C for 15 s and 60 °C for 1 min. For a relative quantitation, the results were normalized to 18S rRNA. Each sample was run in duplicate. The sequence of each primer pair is listed in **Supplemental Table 2.**

### ScRNA-Seq, Library Preparation, Alignment, and Analysis

The scRNA-seq analysis was performed on PLNs or islets from treated mice. PLNs were dissected from the mice, minced into small pieces, and finally digested into single cells using collagenase. Then, they were filtered through a cell strainer to remove tissue debris. Samples from 3-8 individual mice were pooled together for analysis. Islets were isolated from the pancreas using the standard method and then dissociated into single cells by Accutase (Stemcells Technology) digestion for 10 min at 37 °C. The PLNs and islet cells were loaded onto the Chromium Controller (10X Genomics). The resulting samples were processed using the Chromium Single Cell 3′ Library & Gel Bead Kit (10X Genomics, v2) following the manufacturer’s protocol. The libraries were sequenced on Illumina NovaSeq6000.

Raw base call (BCL) files were analyzed using CellRanger (v7.0.0) ^84^. The “fast” command was used to generate FASTQ files, and the “count” command was used to generate raw gene–cell expression matrices. Ambient RNA contamination was inferred and removed using CellBender (v0.2.0) with standard parameters. Mouse genome mm10 was used for the alignment and gencode.vM25 was used for gene annotation and coordinates ^85^. Data from PLN and Islet were analyzed individually. Samples were combined in R using the “Read10X” function from the Seurat package(v4.3.0) ^86^, and an integrated Seurat object was generated. Filtering was conducted by retaining cells with unique molecular identifiers (UMIs) of less than 30,000, none more significant than 500, and mitochondrial content of less than 20 percent. Cell cycle analysis was conducted using CellCycleScoring with a list of cell cycle markers ^87^. Doublets were removed using scDblFinder (v1.12.0) ^88^. The SCTransform workflow was used for count normalization and initial integration and to identify highly variable genes ^89^ using 30 principal components with a resolution of 0.3 for Louvain clustering and UMAP. Cluster marker genes were identified using FindAllMarkers using the Wilcoxon Rank Sum test with the standard parameters. Cell annotation was performed using two approaches: 1) scType, an ultrafast unsupervised method for cell type annotations ^90^, and 2) Manual curation by gene markers to reflect the prediction results.

### Identification of differentially expressed genes (DEGs)

Genes differentially expressed were calculated between the Control and AAT-MSC groups. The R package LIBRA (v1.0.0) was used to perform zero-inflated regression analysis ^91^. Genes were defined as significantly differentially expressed at Benjamini–Hochberg correction FDR < 0.05 and abs (log2(Fold Change)) > 0.3.

### Gene ontology analyses

The functional annotation of the identified DEGs was performed using enrichGO from the clusterProfiler R package ^92^.

### Cell-Cell Interaction analysis

Intercellular communication network analysis was performed using the standard workflow of the R package CellChat (v1.4.0) ^93^.

### Multiplex analysis of cytokine production

Cells from PLNs were cultured in 96-well plates coated with 10 µg/ml anti-CD3 antibody for 96 hours at 37 °C at 5% CO2. The supernatants were then collected and stored at -80 °C. Cytokine levels were measured by analyzing the supernatant using a Premixed Analyte Kit (Mouse Custom 6-Plex, AssayGenie, Ireland), which included TNF-α, IL-1β, IL-17A, IFN-γ, IL-6, and IL-10. The assay was performed following the manufacturer’s instructions, and fluorescence signals were acquired using a CytoFLEX LX flow cytometer (Beckman Coulter Life Sciences, IN, USA).

### Flow cytometry analysis

Spleen and PLNs were isolated from treated mice using standard methods, and the islets were dispersed into single-cell suspension using Accutase (Stemcells Technology) for 10 min at 37 °C. The PLN and spleen cells were fixed with Fixation/Permeabilization Concentrate and Diluent buffer set (Invitrogen, cat. 00-5123-43, 00-5223-56) for 30 minutes on ice. Fixed cells were washed with a flow cytometry staining buffer (FACS buffer, Invitrogen, cat. 00-4222-26). For detection of Tregs, cells were incubated with Brilliant^TM^ violet 605 (BV 605) anti-mouse CD4 (Biolegend, cat. 100547), Phycoerythrin-cyanine7 (PE-Cy7) anti-mouse CD25 (Invitrogen, cat. 25-0251-82), Phycoerythrin (PE) anti-mouse Foxp3 (Invitrogen, cat. 12-5773-82), Alexa fluor 647 (AF647) anti-mouse Helios (BD Biosciences, cat. 563951) and Phycoerythrin-cyanine5 (PE-Cy5) anti-mouse CD152 (CTLA4; Biolegend, cat. 106338) antibodies in permeabilization buffer (Invitrogen, cat. 00-8333-56) at room temperature (RT) for 30 min. For intracellular cytokines detection, cells were cultured in RPMI 1640 media (10% fetal bovine serum, 200 μg/ml penicillin, and 50 uM 2-mercaptoethanol) containing phorbol 12-myristate 13-acetate (PMA; 50 ng/ml, Sigma-Aldrich, cat. P8139), ionomycin (1 ug/ml; Sigma-Aldrich, cat. I9657) and 1x protein transport inhibitor (BD bioscience, cat. 555029) for 4 h to ensure intracellular accumulation of cytokines. After stimulation, cells were washed with FACS buffer and fixation. Fixed cells were incubated with BV 605 anti-mouse CD4, PE-Cy7 anti-mouse CD25, AF647 anti-mouse IFN-γ (Biolegend, cat. 505814) and Peridinin chlorophyll-cyanine5.5 (PerCP-Cy5.5) anti-mouse IL-17A (Biolegend, cat. 506920) in permeabilization buffer at RT for 30 min. For exhausted CD8^+^ T cells detection, fixed cells were incubated with Allophycocyanin-cyanine7 (APC-Cy7) anti-mouse CD8 (Biolegend, cat. 100714), Brilliant™ Violet 421 (BV 421) anti-mouse CD279 (PD-1; Biolegend, cat. 135221), PE anti-mouse TCF7/TCF1 (BD bioscience, cat. 564217), PerCP-Cy5.5 anti-mouse CD366 (Tim-3; Biolegend, cat. 119718), AF647 anti-mouse Tox (BD bioscience, cat. 568356) and PE-Cy7 anti-mouse TIGIT (Biolegend, cat. 142108) in permeabilization buffer at RT for 30 min. Flow cytometry analysis was performed on BD LSRFortessa Cell Analyzer (BD Biosciences) and analyzed using FlowJo software. A list of antibodies used can be found in **Supplemental Table 1.**

### T cell immunosuppressive assay

PLNs were isolated from Foxp3^GFP^NOD mice three weeks after infusion with AAT-MSCs. CD4^+^CD25^+^GFP^+^ Treg cells were sorted from the PLNs of mice infused with either AAT-MSCs or a control (CTR) treatment. T cells were isolated from splenocytes of C57BL/6 mice by flow cytometry to serve as responder cells. The sorted T cells were labeled with the CTV kit (Invitrogen, cat. C34557) for 10 minutes at 37°C. Treg cells (3×10^5) were then co-cultured with the labeled T cells at a 1:1 ratio in RPMI 1640 medium supplemented with 10% FBS, 2-Mercaptoethanol (1:1000 dilution), and glutamine (Thermo Fisher) in 96-well plates. Cells were stimulated with anti-CD3 (2 µg/mL; BioXCell, cat. BE0002) and anti-CD28 (2 µg/mL; BioXCell, cat. BE0015-1) antibodies for 3 days. T-cell proliferation was assessed by flow cytometry using the standard method ^94^.

### Co-culture of AAT-MSCs with human PBMCs

AAT-MSCs (3×10^5^) were seeded in 96 well plates and cultured in a 5% CO2 incubator for 2 hours at 37°C. Human PBMC (3×10^5^ per well) was added in each well for co-culture. After 4 days, the PBMCs were collected and stained with APC anti-human CD4 (BD Biosciences, cat. 561840), PerCP-Cy5.5 anti-human CD25 (BD Biosciences, cat. 560503), and Alexa fluor 700 (AF700) anti-human CD127 (Biolegend, cat. 351343) antibodies. Numbers of CD4^+^CD25^high^CD127^lo^ proliferation were analyzed by flow cytometry.

### Co-culture of Tregs, AAT-MSCs with human CD8^+^ T cells and islets

CD4^+^CD25^+^GFP^+^ Tregs and GFP^-^T cells were isolated from FoxP3^EGFP^ NOD mice and stimulated with anti-CD3 (5 µg/ml) and anti-CD28 (5 µg/ml) antibodies. Cells were then co-cultured/”educated” with AAT-MSCs at a 1:1 ratio for 72 hours. Subsequently, the educated Tregs or non-Tregs were co-cultured with CD8^+^ T cells for 72 hours, and the presence of exhausted CD8^+^ T cells was quantified by flow cytometry. CD8^+^ T cells were then separated from other cells using the MagniSort^TM^ Mouse CD8 T cells Enrichment Kit (Invitrogen, Waltham, MA) and further co-cultured with mouse islets for 3 days before cell death analysis in the islets.

### Islet live and death and Terminal deoxynucleotidyl transferase dUTP Nick End Labeling (TUNEL) staining

For live and cell staining, fresh islets were incubated with 0.1 µM SYTO-13 and 10 μg/ml EB in PBS for 15 min in the dark. The stained islets were then observed under a fluorescent microscope. For apoptosis detection using the TUNEL assay, formalin-fixed, cryopreserved islet sections were washed with PBS for 5 min. the TUNEL reaction was performed using the In Situ Cell Death Detection Kit, Fluorescein (Roche) for 1 hour at room temperature. After washing the section with PBS, slide sections were treated with 0.1% Triton-X 100 for 15 min at room temperature to permeablize the cells. The sections were then incubated overnight at 4° C with an anti-insulin antibody. The following day, the sections were washed again and incubated with 2nd antibody at RT for one hour. Fluorescence images were captured using an EVOS M5000 fluorescence microscope (Thermo Fisher).

### Statistical Analysis

Data are expressed as mean ± standard deviation (SD) or standard error of the mean (SEM). Z scores were calculated for some variables. Differences among groups were tested by one-way ANOVA with post hoc analysis. Comparisons between the two groups were evaluated using Student’s t-test. A value of p < 0.05 was considered significantly different.

## ACKNOWLEDGMENT

This study was supported in part by the National Institutes of Health (DK105183, DK 120394, DK118529, and DK125464) and the Department of Veterans Affairs (VA-ORD BLR&D Merit I01BX004536). Some illustrations were created in BioRender. Wang, H. (2025) https://BioRender.com/elzcms9

## DATA AVAILABILITY

The datasets generated and/or analyzed in this study are available from the corresponding author by a written request.

