## Supplemental data for "Alpha-1 Antitrypsin Overexpressing Mesenchymal Stem/Stromal Cells Reverses Type 1 Diabetes via Promoting Treg Function and CD8^+^ T cell exhaustion"

#### Supplement Fig S1

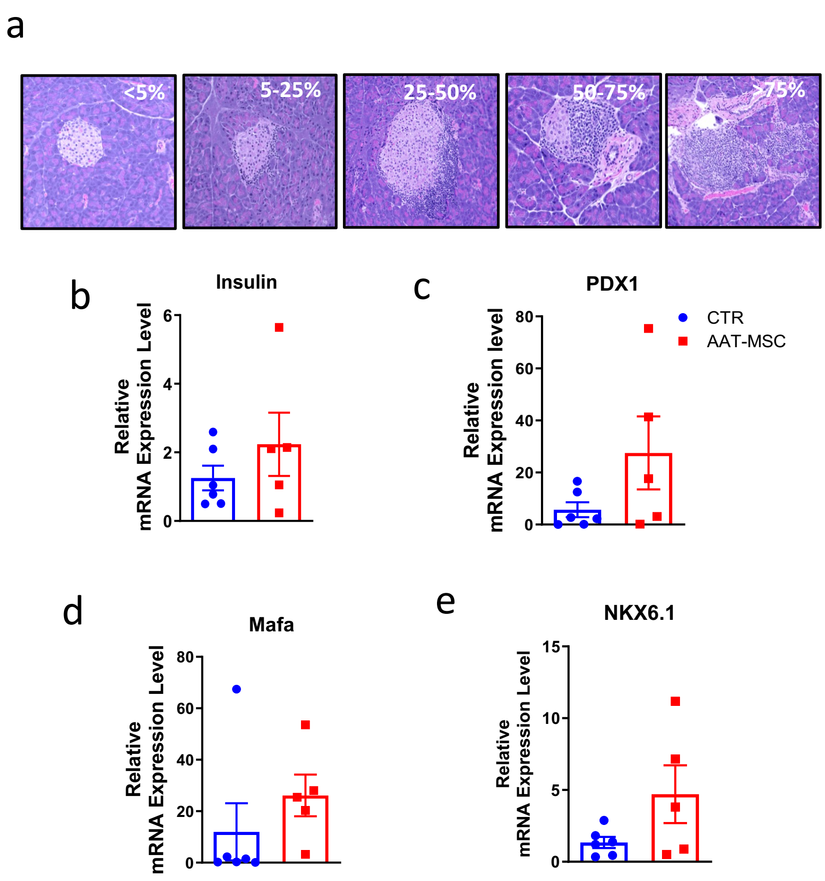


**Supplemental Fig. S1. Islets from AAT-MSC-treated mice exhibit a trend of increased mRNA expression of key transcriptional factors essential for islet function. a:** Hematoxylin and Eosin (H.E.) staining of pancreatic tissues reveals the morphology of islets with varying degrees of immune cell infiltration. PCR analysis of **(b)** PDX1, **(c)** Insulin, **(d)** Mafa, and **(e)** NKX6.1 in islets from control or AAT-MSC treated mice. Each dot represents a sample from an individual mouse.

### Supplemental Fig S2.

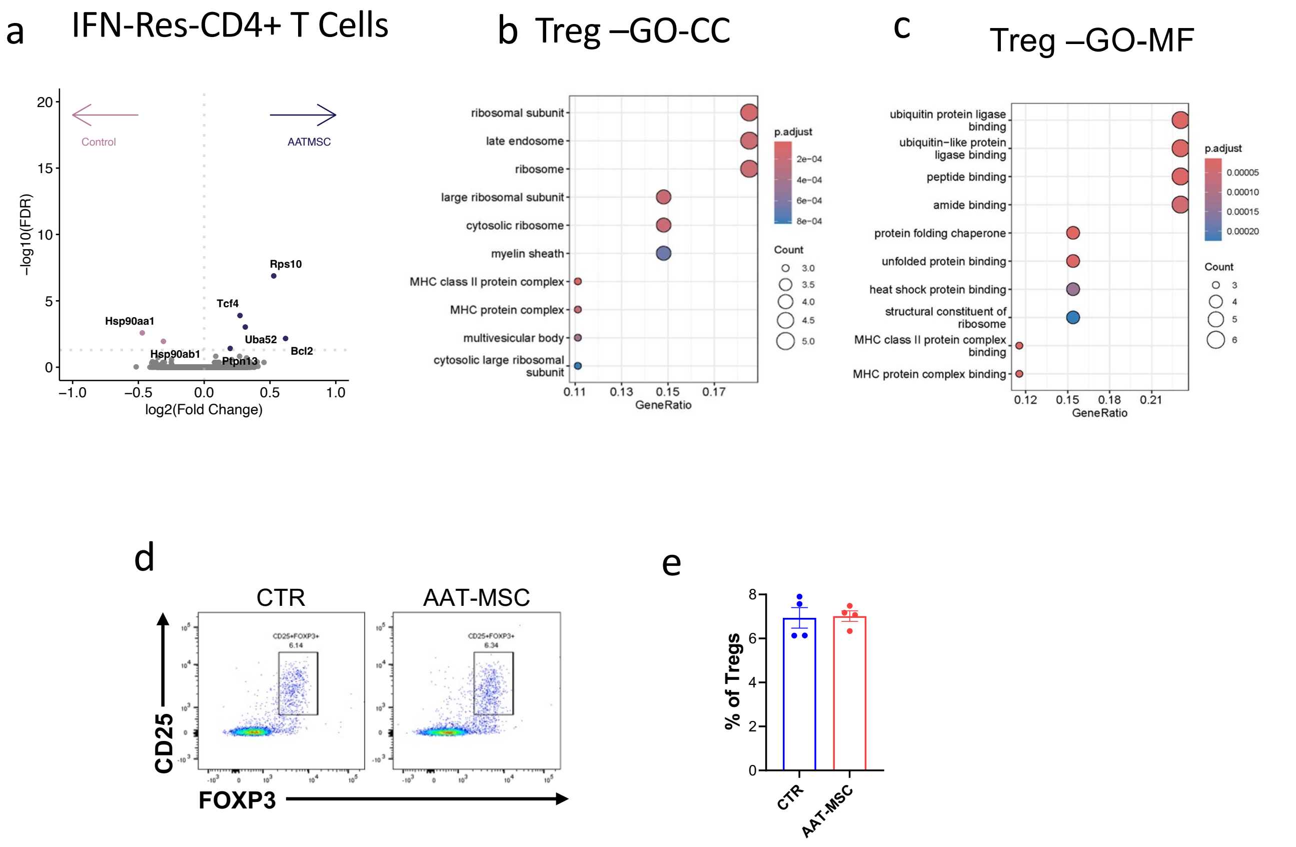


**Supplemental Fig S2.** Additional characterization of $CD4^+$ T cell populations in the PLNs of CTR and AAT-MSC-treated mice. **a.** Differentially expressed genes (DEGs) from IFN-responsive $CD4^+$ T cells **(b),** GO pathway analysis for cellular component (CC) and molecular function (MF) in Tregs from CTR and AAT-MSC-treated mice. **d, e**: Impact of AAT-MSC therapy on Treg numbers three weeks after AAT-MSC or control infusion measured by flow cytometry (**d**) and quantified as % of Treg among total $CD4^+$ T cells (**e**). n=4 per group.

#### Supplemental Fig S3

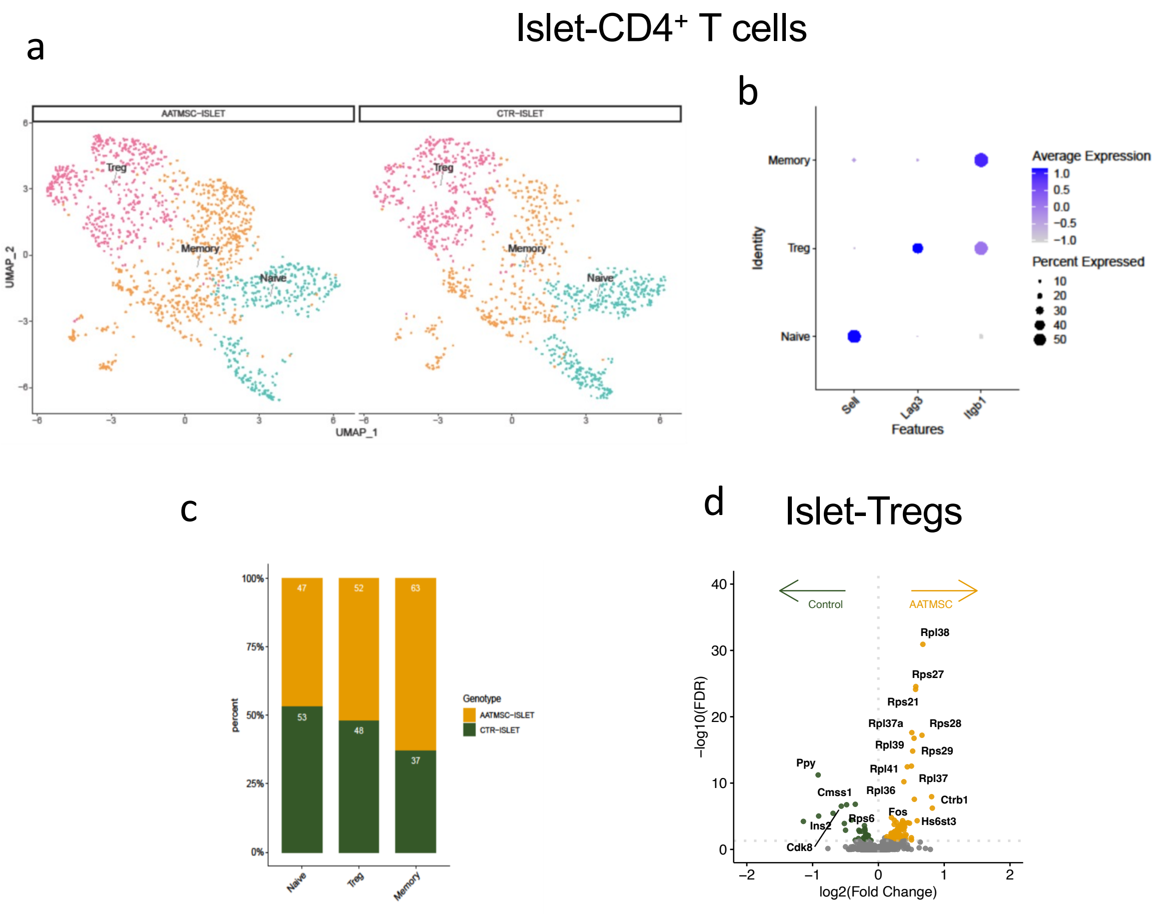


**Supplemental Fig. S3**. **a**. t-SNE projection plot of CD4$^+$ T cells in the islets from CTR or AAT-MSC-treated mice. **b**. GSEA summary of gene signature for each cell subpopulation. **c.** Subtype distribution of CD4$^+$ T cells in the islets of the CTR and AAT-MSC-treated mice. **d**. DEGs of islet Tregs from AAT-MSC-treated and control mice.

#### Supplemental Fig S4

PLN-CD8$^+$ T cells

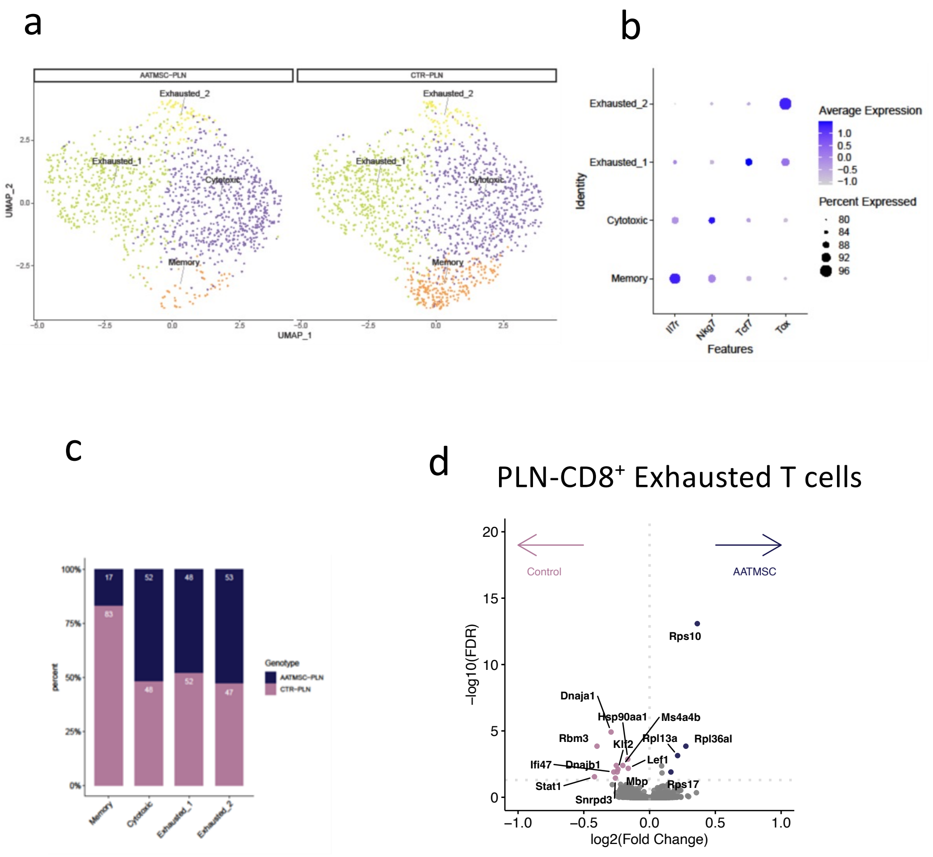


**Supplemental Fig. S4. CD8$^+$ T cell populations in islets from AAT-treated or control mice. a**. t-SNE projection plot of CD8$^+$ T cells in the PLNs from the mice treated with AAT-MSC and CTR. **b**. GSEA summary of gene signature for each population. **c.** Subtype distribution of CD4$^+$ T cells in PLNs of CTR and AAT-MSC-treated mice. **d**. Volcano plots showing significantly changed gene expression in exhausted CD8$^+$ T cells from islets of CTR and AAT-MSC-treated mice.

#### Supplemental Fig. S5

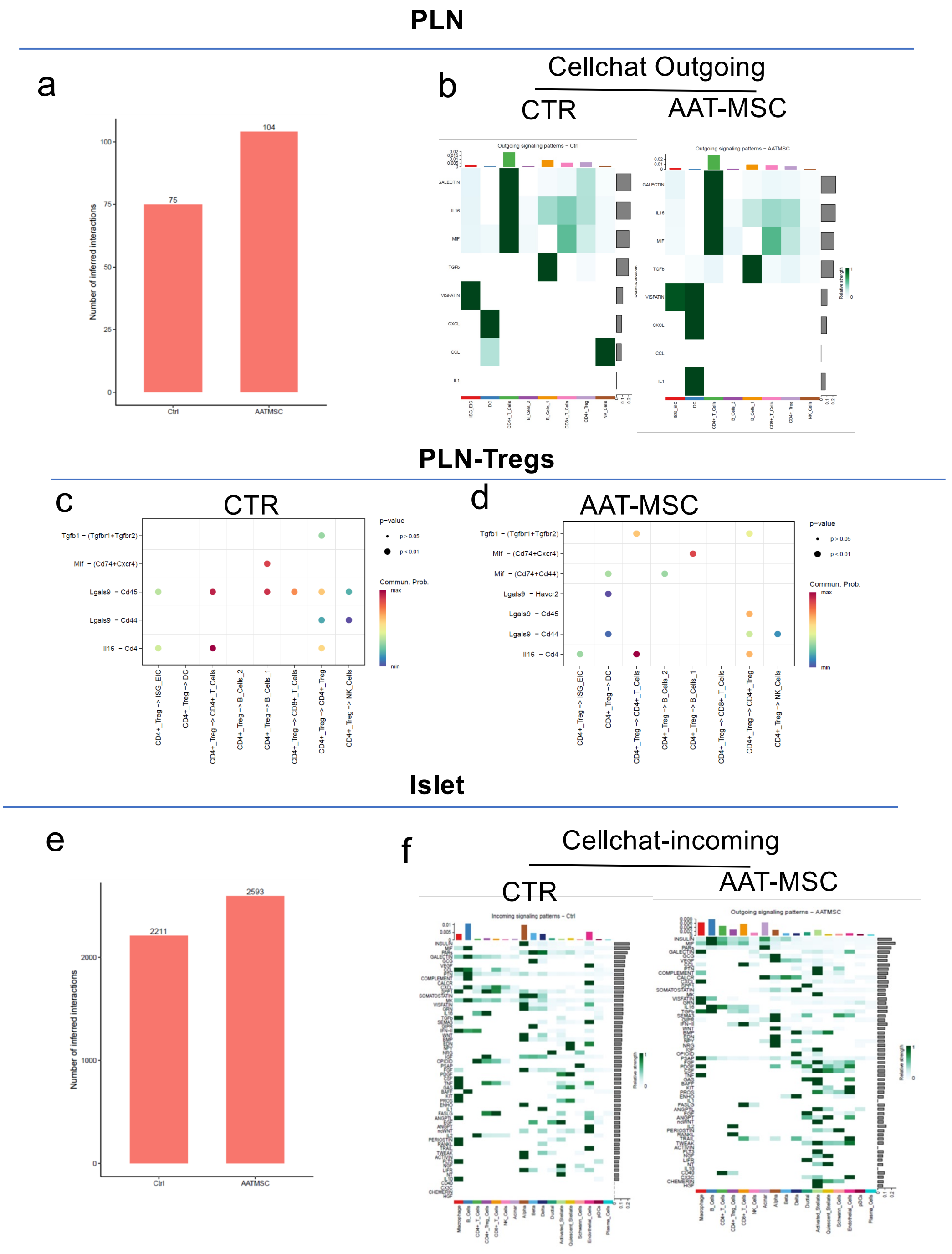


**Supplemental Fig. S5. a.** Numbers of inferred cell interactions from PLNs from control or AAT-treated mice. **b**. Heatmaps of outgoing signals in PLN cells from CTR or AAT-MSC-treated mice using Cellchat analysis. **c &d.** Dot plots of ligand-receptor prediction analysis between PLN-Tregs and other immune cells from CTR or AAT-MSC-treated mice. **e.** Numbers of inferred interactions in cells from islets from control or AAT-treated mice. **b**. Heatmaps of incoming signals in PLN cells from CTR or AAT-MSC-treated mice done with Cellchat.

#### Supplemental Fig. S6

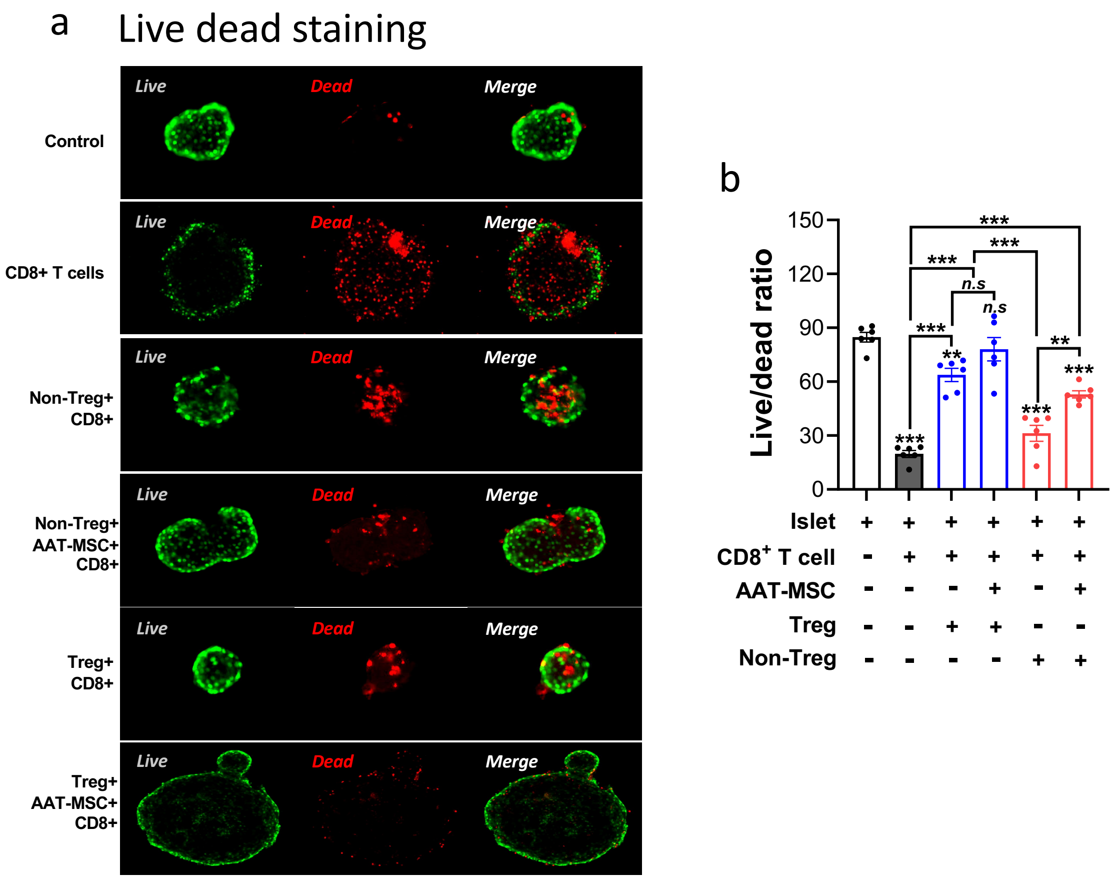


**Supplemental Fig. S6. Fluorescence micrograph of islets stained with SYTO-13 and ethidium bromide (EB). a.** Fluorescent microscope images of Live and death staining and quantification (**b**) of islet cells were performed in the presence of $CD8^+$ T cells, with Tregs or non-Tregs, and/or AAT-MSC co-culture, followed by staining with SYTO-13 and EB. Red: dead cells, green: live cells. $*p < 0.05$, $**p < 0.01$. $***p < 0.001$. Each group was compared with islet alone or islet with $CD8^+$ T cell co-culture group. One-way ANOVA with post-hoc correction.

### Supplement Fig. S7

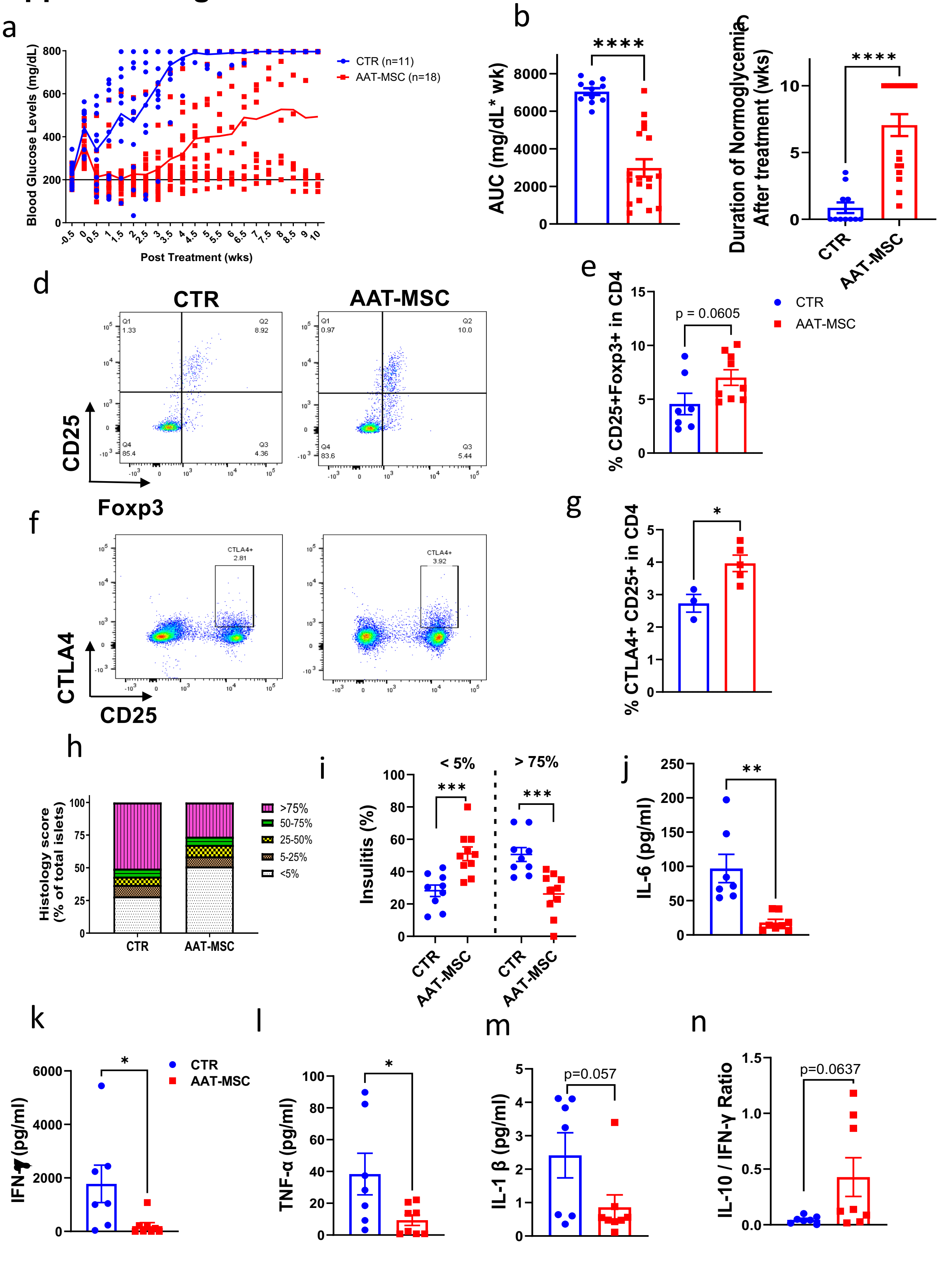

**Supplemental Fig. S7. AAT-MSC treatment reverses diabetes in mice with new-onset T1D.** a. Non-fasting blood glucose levels of new-onset T1D mice without treatment (CTR, n=11) or treated with AAT-MSC cells (1 x $10^6$/mouse, i.v. infusion), once blood glucose level consistently exceeds 250 mg/dL (AAT-MSC, n=18). **b**. The AUC of blood glucose levels during the following period was calculated in each mouse. **c**. Duration of normoglycemia post-treatment in T1D mice. **d. e. f. g**. Analysis of Treg population in PLNs of T1D mice 4 weeks post-treatment. Scatter plots (**d, f**) and the percentage of $CD4^+CD25^+FOXP3^+$ cells and $CD4^+CD25^+CTLA4^+$ cells in $CD4^+$ (e, g) were significantly higher in AAT-MSC-treated mice. **d and** e: n=7 in CTR and n=9 in AAT-MSC. **f and g**: n=3 in CTR and n=5 in AAT-MSC. Each dot presents one mouse. **h, i and j.** Evaluation of insulitis in pancreatic tissue sections from control or AAT-MSC-treated T1D mice, stained with H.E. **i.** Individual histology scores were determined from 588 islets in the control group and 493 islets in the AAT-MSC-treated group. **J-n**. Measurement of IL-6, IFN-γ, TNF-α, IL-1,β and the ratio of IL-10/IFN-γ from anti-CD3 stimulated PLN cells of CTR or AAT-treated mice. Data are presented as mean ± SEM. $*P < 0.05$, $**P < 0.01$, determined by unpaired two-tailed t-test with F test to compare variances.

**Table S1. Antibodies used in the study.**

| Antibody Name | Type | Dilution | Source | Identifier |
|---|---|---|---|---|
| CD4 | Rat, monoclonal | 1:100 | BioLegend | Cat# 100547; RRID:AB_11125962 |
| CD25 (PC61.5) | Rat, monoclonal | 1:100 | Invitrogen | Cat# 25-0251-82; RRID:AB_469608 |
| FOXP3 (FJK-16s) | Rat, monoclonal | 1:100 | Invitrogen | Cat# 12-5773-82; RRID:AB_465936 |
| Helios | Armenian hamster, monoclonal | 1:100 | BD Biosciences | Cat# 563951; RRID:AB_2738506 |
| CD152 | Armenian hamster, monoclonal | 1:100 | BioLegend | Cat# 106338; RRID:AB_3083256 |
| IFN-γ | Rat, monoclonal | 1:100 | BioLegend | Cat# 505814; RRID:AB_493314 |
| IL-17A | Rat, monoclonal | 1:100 | BioLegend | Cat# 506920; RRID:AB_961384 |
| CD8a | Rat, monoclonal | 1:100 | BioLegend | Cat# 100714; RRID:AB_312752 |
| CD279 (PD-1) | Rat, monoclonal | 1:100 | BioLegend | Cat# 135221; RRID:AB_2561447 |
| TCF-7/TCF-1 | Mouse, monoclonal | 1:100 | BD Biosciences | Cat# 564217; RRID:AB_2687845 |
| CD366 (Tim-3) | Rat, monoclonal | 1:100 | BioLegend | Cat# 119718; RRID:AB_2571934 |
| TOX | Rat, monoclonal | 1:100 | BD Biosciences | Cat# 568356 |
| TIGIT (Vstm3) | Mouse, monoclonal | 1:100 | BioLegend | Cat# 142108; RRID:AB_2565648 |
| CD3 | Rat, monoclonal | 2 µg/mL | Bio X Cell | Cat# BE0002; RRID:AB_1107630 |
| CD28 | Syrian hamster, monoclonal | 2 µg/mL | Bio X Cell | Cat# BE0015-1; RRID:AB_1107624 |
| CD4 | Mouse, monoclonal | 1:100 | BD Biosciences | Cat# 561840; RRID:AB_398593 |
| CD25 | Mouse, monoclonal | 1:100 | BD Biosciences | Cat# 56050; RRID:AB_1727453 |
| CD127 (IL-7Rα) | Mouse, monoclonal | 1:100 | BioLegend | Cat# 351343; RRID:AB_2566199 |

**Table S2. Primer sequences for targeted genes.**

| Target | Forward primer 5'-3' | Reverse primer 5'-3' |
|---|---|---|
| PDX1 | ACTTAACCTAGGCGTCGCACAAGA | GGCATCAGAAGCAGCCTCAAAGTT |
| Ins1 | ATCCACAATGCCACGCTTCT | AAACCCACCCAGGCTTTTG |
| Mafa | AGGAGGAGGTCATCCGACTG | CTTCTCGCTCTCCAGAATGTG |
| NKX6.1 | TCTGGACAGCAAATCTTCGCCC | ACTTGGTCCTGCGGTTCTGGAA |
